# Autism risk gene POGZ promotes chromatin accessibility and expression of clustered synaptic genes

**DOI:** 10.1101/2021.04.07.438852

**Authors:** Eirene Markenscoff-Papadimitriou, Fadya Binyameen, Sean Whalen, James Price, Kenneth Lim, Rinaldo Catta-Preta, Emily Ling-Lin Pai, Xin Mu, Duan Xu, Katherine S. Pollard, Alex Nord, Matthew W. State, John L. Rubenstein

## Abstract

*De novo* mutations in *POGZ*, which encodes the chromatin regulator *Pogo* Transposable Element with ZNF Domain protein, are strongly associated with autism spectrum disorder (ASD). Here we find that in the developing mouse and human brain POGZ binds predominantly euchromatic loci and these are enriched for human neurodevelopmental disorder genes and transposable elements. We profile chromatin accessibility and gene expression in *Pogz^−/−^* mice and find that POGZ promotes chromatin accessibility of candidate regulatory elements (REs) and the expression of clustered synaptic genes. We further demonstrate that POGZ forms a nuclear complex and co-occupies loci with HP1γ and ADNP, another high-confidence ASD risk gene. In *Pogz*^+/−^ mice, *Adnp* expression is reduced. We postulate that reduced POGZ dosage disrupts cortical function through alterations in the POGZ-ADNP balance which modifies neuronal gene expression.

## Introduction

Chromatin packaging of DNA is a dynamic process that determines the transcriptional potential of a cell. Chromatin regulators compartmentalize chromatin into transcriptionally active (euchromatic) or silent (heterochromatic) domains by remodeling nucleosomes and modifying the histone code. Euchromatin is characterized by accessible DNA, while heterochromatin is characterized by condensed chromatin and H3K9me3 and H4K20me3 histone modifications. Chromatin regulators prepare the underlying chromatin landscape on which transcription factor programs that determine cell subtype specification are executed with spatiotemporal precision (Nord et al. 2015). Mutations in chromatin genes are strongly linked to autism spectrum disorders (Satterstrom et al. 2020; Sanders et al. 2015; De Rubeis et al. 2014) and a range of other neurodevelopmental phenotypes (Deciphering Developmental Disorders Study 2015). Understanding how mutations in chromatin regulators impact gene regulation during mouse brain development can illuminate the underlying mechanisms of social and intellectual disability and potentially other neuropsychiatric syndromes (Gompers et al. 2017; Jung et al. 2017; Cappi et al. 2020).

*POGZ* is a high confidence ASD risk gene identified by whole exome sequencing of patient cohorts (Satterstrom et al. 2020; Sanders et al. 2015; De Rubeis et al. 2014; Stessman et al. 2016; Iossifov et al. 2014). Subsequent to its identification as a large-effect risk gene in studies of individuals with idiopathic ASD, “genotype-first” analyses led to the characterization of White-Sutton syndrome, defined by mutations in POGZ. This syndrome is marked by distinctive facial features along with varying degrees of intellectual disability (ID), ASD, neurological and gastrointestinal abnormalities (Batzir et al. 2020).

*POGZ* encodes a protein whose function is not well understood. *In silico* analysis suggests that it is a DNA binding protein with N terminal zinc finger (ZNF) domains and a C terminal DNA binding and transposase domain. ASD associated mutations disrupt POGZ’s DNA binding activity (Matsumura et al. 2016) and reduce neurite outgrowth *in vitro* (Hashimoto et al. 2016; Zhao et al. 2019). POGZ is expressed in the developing brain and POGZ knockdown and knockout leads to impairments in mouse cortical development and alteration in social behaviors (Matsumura et al. 2020; Suliman-Lavie et al. 2020). We have previously analyzed POGZ heterozygote mice and found they exhibit anxiety-related avoidance behavior and reduced inhibition in the medial prefrontal cortex (Cunniff et al. 2020). Thus, *POGZ* is a *bona fide* neurodevelopmental disorder (NDD) gene that dramatically increases the risk in humans for ASD and intellectual disability, which can be recapitulated in mouse models.

Mechanistic insight into the neurodevelopmental function of POGZ is lacking. A previous publication reported that POGZ binds heterochromatin proteins through its ZNF domains (Nozawa et al. 2010); as a result, POGZ has been thought to function as a repressor of transcription (Suliman-Lavie et al. 2020). Here, we analyzed transcriptional and epigenetic changes in a *Pogz* constitutive null mouse, as well as POGZ binding in mouse and human cortical tissues. Our analysis suggests that POGZ acts both as a transcriptional activator and a repressor. We find POGZ binds euchromatic loci with Heterochromatin Protein 1γ (HP1γ), a component of euchromatin, and ADNP, another confirmed ASD risk gene (Sanders et al 2015, Satterstrom et al 2020). Our results provide a mechanistic dissection of POGZ’s molecular function in embryonic forebrain development and elucidate direct targets of POGZ.

## Results

### Generation of POGZ constitutive null

POGZ is expressed prenatally in the mouse cortex and ganglionic eminences (LGE and MGE; basal ganglia); *in situ* hybridization (ISH) of *Pogz* across developmental stages indicates decreasing expression in postnatal brain (Figure S1A-C). In the developing cortex *Pogz* is broadly expressed from the ventricular zone to the cortical plate, thus spanning neuronal progenitors and newborn neurons, and is more strongly expressed in neurons (Figure 1C).

**Figure 1).**
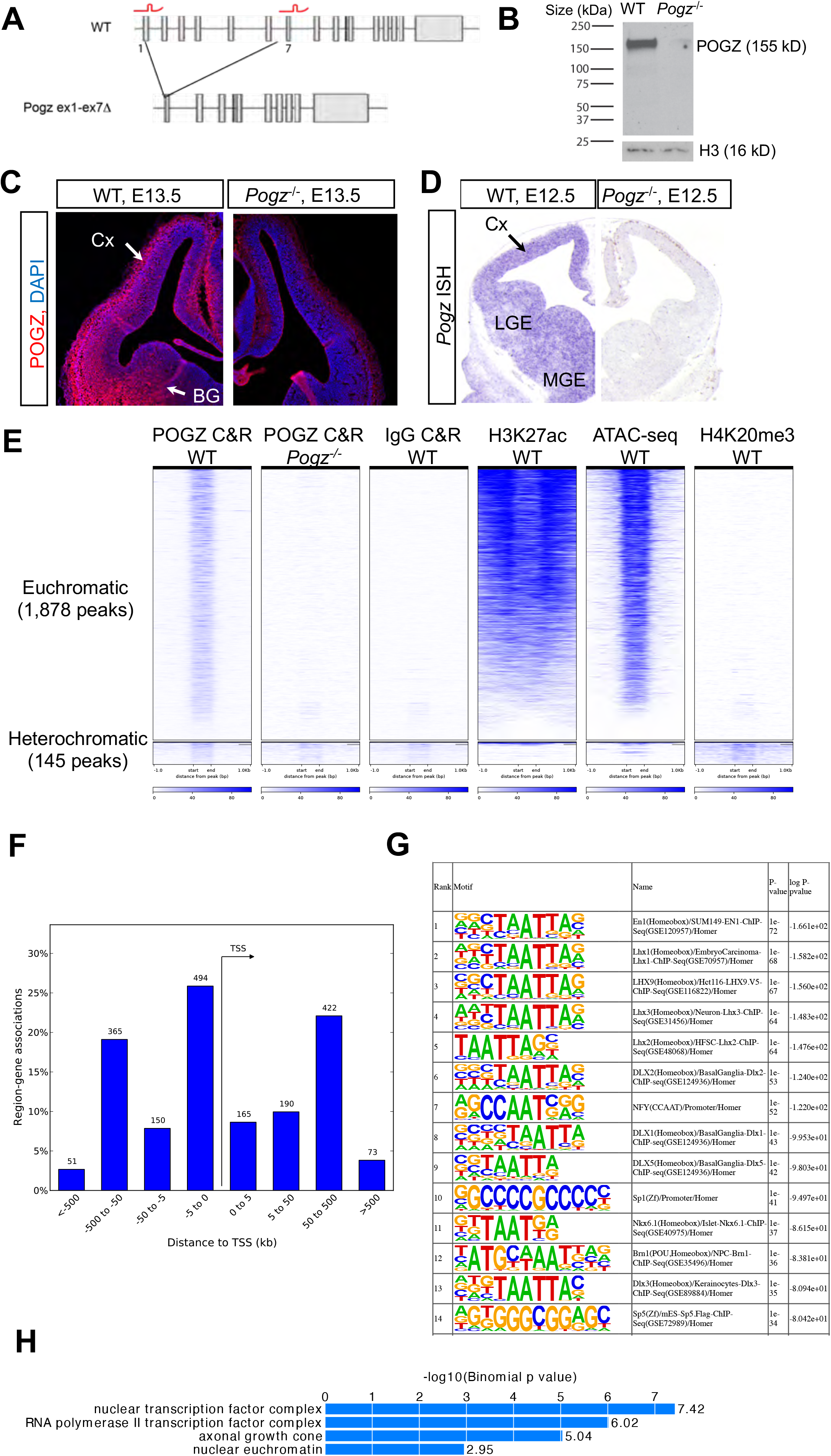
POGZ binds euchromatic loci in the developing mouse telencephalon. 1A) CRISPR-Cas9 generation of POGZ knockout allele. sgRNAs (red) targeting exons 1 and 7 of mouse *Pogz* are indicated. Resulting 10.5 kilobase deletion of *Pogz^−/−^* allele is shown. 1B) Western blot using anti-POGZ antibody in nuclear extracts from wildtype and *Pogz^−/−^* mouse cortex, E13.5. Histone H3 loading control. 1C) Anti-POGZ immunostaining (red) and DAPI (blue) in E13.5 WT and *Pogz^−/−^* telencephalon sections. POGZ expression in basal ganglia (BG) and cortex (Cx) is indicated. 1D) ISH of E12.5 WT and *Pogz^−/−^* telencephalon sections with *Pogz* antisense probe (purple). *Pogz* expression in MGE, LGE, and cortex is indicated. 1E) Heat map of POGZ C&R reads across consensus peaks (*n* = 2,023) in the mouse genome, in E13.5 telencephalon. Each row represents a 2 kb window centered on peak midpoints, sorted by the POGZ C&R signal. *Pogz^−/−^* and IgG controls are shown. ATAC-seq, H3K27ac, and H4K20me3 ChIP-seq signal from E13.5 wildtype shown to the right. Euchromatic peaks on the top row, heterochromatic peaks on the bottom row. RPM, reads per million. 1F) C&R peaks linked to nearest genes, binned by distance and orientation from gene TSS using GREAT. 1G) HOMER motif analysis of POGZ C&R sequences. 1H) GO terms significantly enriched in POGZ C&R nearest genes.

We generated a constitutive *Pogz* deletion allele by pronuclear injection of fertilized murine C57/Bl6 eggs with CRISPR-Cas9 protein and guide RNAs targeting exons 1 and 6, a 10 kb span (Figure 1A). Founders were screened by PCR (see Methods). One founder contained a 10 kb deletion which generated a premature stop codon (Figure S1D). We out-crossed this founder to C57/Bl6 mice for ten generations. *Pogz^−/−^* mice die at embryonic day E15.5, as previously reported, of uncertain etiology (Gudmundsdottir et al. 2018); *Pogz^+/−^* mice survive and are fertile. At E13.5 we observed no *Pogz* expression in homozygous knockouts based on Western blot, immunohistochemistry and ISH (Figure 1B, 1C, 1D).

Reduced cortical neurogenesis has been described in *Pogz* shRNA knockdown experiments (Matsumura et al. 2020). Thus, we analyzed *Pogz^−/−^* cortex at E13.5 for markers of proliferation and neurogenesis. Immunostaining for mitotic marker Phosphohistone H3 (PHH3) showed a modest increase in PHH3+ cells in the *Pogz^−/−^* ventricular zone (Figure S2A), however the trend was not significant. There was no significant change in the thickness of the TBR2^+^ sub-ventricular zone or the β-III Tubulin^+^ cortical plate in *Pogz^−/−^* (Figures S2B,C). ISH for Layer 5 and Layer 6 neuron markers *Fezf2* and *Tbr1*, respectively, showed no difference in cortical neuron fate or lamination (Figures S2D, S2E). Thus, we conclude the E13.5 *Pogz^−/−^* cortex does not have a strong neurogenesis phenotype.

### POGZ occupies euchromatin loci

To dissect the molecular function of POGZ, we used *Pogz^−/−^* mice at E13.5. Immunofluorescence (IF) of POGZ in E13.5 medial ganglionic eminence shows nuclear staining that appears euchromatic as it is not restricted to DAPI positive heterochromatin (Figure S1E). For an unbiased screen of POGZ occupancy genome-wide in embryonic nuclei, we performed CUT&RUN (C&R) (Skene, Henikoff, and Henikoff 2018) in E13.5 dissected wild type telencephalons using anti-POGZ antibody. *Pogz^−/−^* embryos and IgG were used as negative controls. We identified 2,023 consensus POGZ peaks in C&R analysis of duplicate experiments (see Methods, Table S1). POGZ C&R peaks have greatly reduced signal in *Pogz^−/−^* and wildtype IgG experiments, demonstrating the validity of POGZ interactions (Figure 1E).

To establish whether POGZ occupies euchromatic or heterochromatic loci, we compared POGZ bound loci to ATAC-seq and H3K27ac and H4K20me3 ChIP-seq we generated from wildtype E13.5 cortex. Unexpectedly, because POGZ has been shown to bind heterochromatin protein HP1α (Nozawa et al. 2010), we find POGZ occupancy occurs predominantly at euchromatic rather than heterochromatic loci. 92% of POGZ occupied loci contain accessible chromatin and are enriched for H3K27ac, while 8% overlap peaks for the heterochromatin hallmark H4K20me3 (Figure 1E).

POGZ occupies transcription start sites (29% of peaks) and distal intergenic regions (71%), suggesting it may act as a transcriptional regulator (Figure 1F). 96 validated enhancers that have activity in the E11.5 mouse embryonic brain (Visel et al. 2007) overlap with POGZ C&R peaks, indicating that POGZ binds brain enhancers. Furthermore, HOMER motif analysis of POGZ C&R peaks shows that they are highly enriched for homeobox (e.g. DLX, LHX, POU) and ZNF motifs (e.g. SP), which are found in TFs that bind telencephalic enhancers (Figure 1G) (Lindtner et al. 2019; Sandberg et al. 2016). *De novo* motif discovery analysis identified putative POGZ binding motifs which will be discussed later (Figure 7).

Genes near POGZ-occupied loci are enriched for gene ontology (GO) terms that include “nuclear euchromatin” and “axonal growth cone” (Figure 1H), suggesting that POGZ may transcriptionally regulate genes that encode components of euchromatin and axon growth. Taken together, POGZ C&R, epigenetic analyses, and IF experiments provide evidence that POGZ binds predominantly euchromatic loci and also directly regulates genes associated with euchromatin.

### Dysregulation of neuronal genes in *Pogz^−/−^* embryos

For an unbiased screen of genes dysregulated in *Pogz^−/−^* we performed bulk RNA-seq. We compared gene expression of E13.5 *Pogz^−/−^* and wildtype littermates, and analyzed the cortex in triplicate and the basal ganglia in duplicate. In cortex, 177 genes were significantly down-regulated and 154 were significantly up-regulated (qvalue<0.05) (Figure 2A, see Table S2). In basal ganglia, fifteen genes were down-regulated and sixteen up-regulated (qvalue<0.05) (Figure 2B).

**Figure 2).**
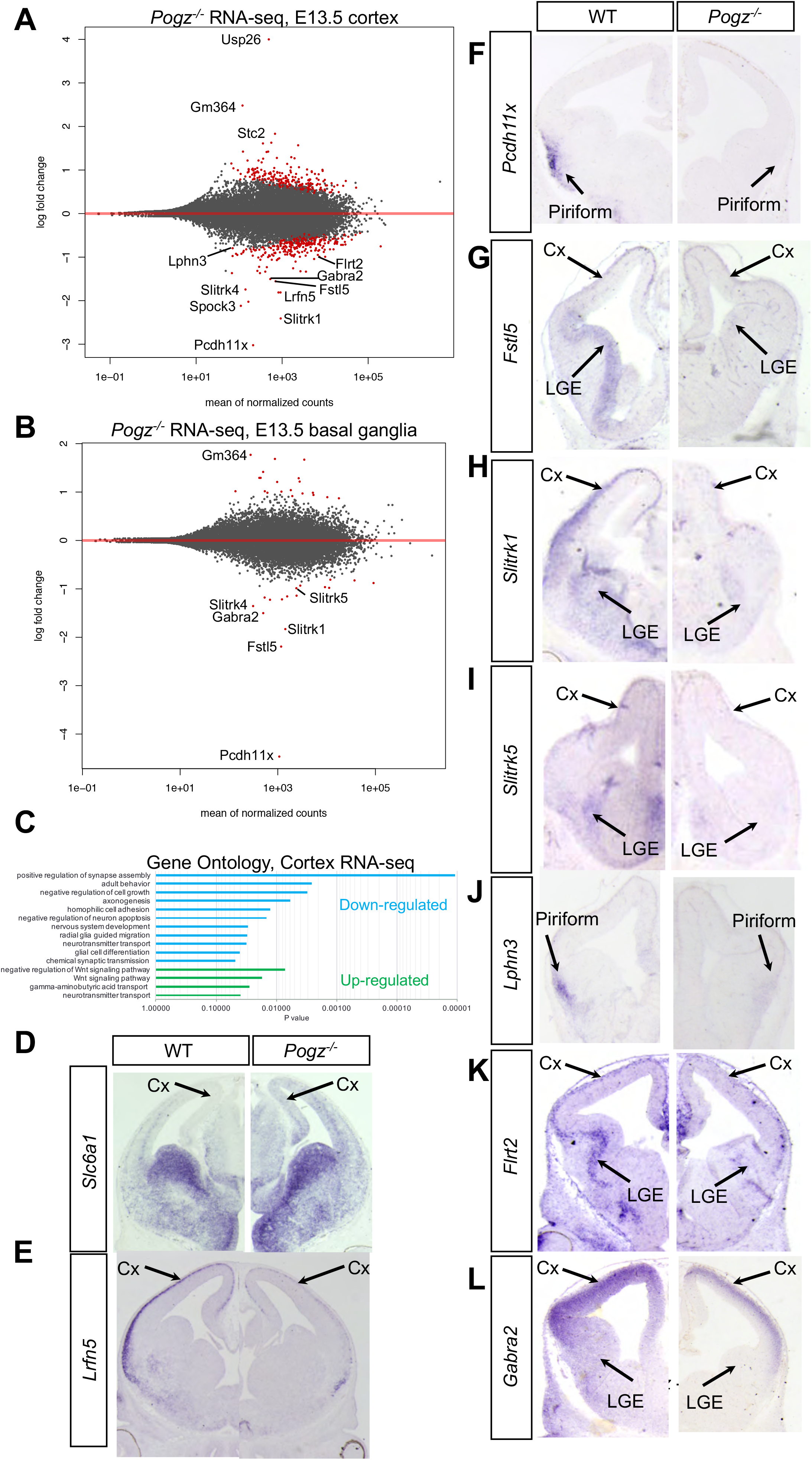
Neuronal genes down-regulated in *Pogz^−/−^* forebrain. 2A,B) MA plots showing differentially expressed genes in E13.5 *Pogz^−/−^* cortex and basal ganglia RNA-seq compared to wildtype controls. Significantly (qvalue<0.05) differentially expressed genes are indicated in red dots. 2D-L) ISH expression and validation of genes up-regulated (D) and down-regulated (E-L) in *Pogz^−/−^* at E13.5.

Down-regulated genes in cortex were enriched for GO terms such as “positive regulation of synapse assembly”, “axonogenesis”, and “homophilic cell adhesion” (Methods, see Figure 2C). Up-regulated genes were enriched for terms such as “Wnt signaling”, “neurotransmitter transport,” and “GABA transport” (Figure 2C). The most highly down-regulated gene in both tissues was *Pcdh11x*, whose expression decreased 8-fold in cortex and 22-fold in basal ganglia.

We explored the most significant (qvalue<0.01) down-regulated genes in *Pogz^−/−^.* Notably, in *Pogz^−/−^* cortex, *Slitrk1,2,4, and 5* were downregulated (Figure S3A), and in basal ganglia *Slitrk1,4*, and 5 were downregulated (Figure S3B). *Slitrk* family genes encode leucine-rich repeat extracellular proteins that promote axon extension and synapse formation (Aruga and Mikoshiba 2003; Abelson et al. 2005; Linhoff et al. 2009). *Slitrk1* is a Tourette’s disorder candidate gene (Abelson et al. 2005; O’Roak et al. 2010) and *Slitrk5* mouse mutants have OCD like phenotypes (Shmelkov et al. 2010). *Lrfn5*, encoding another leucine-rich repeat extracellular protein implicated in NDDs (Cappuccio et al. 2019), was down-regulated in *Pogz^−/−^* as well.

*Lphn3* and *Flrt2* were down-regulated two-fold in *Pogz^−/−^* cortex (Figure 2A). *Lphn3* encodes a GPCR that promotes excitatory synapse formation by trans-synaptic binding of FLRT family and Teneurin proteins (Sando, Jiang, and Südhof 2019). *Gabra2, Gabra4*, and *Gabrg1* encode components of the GABA_A_ receptor and were down-regulated 2.5-fold in *Pogz^−/−^* cortex and basal ganglia (Figure 2A, 2B). Genes encoding GABA transporters SLC6A1 and SLC6A11 were up-regulated in *Pogz^−/−^* cortex (Figure S3C); *Slc6a1* is associated with ASD and myoclonic atonic epilepsy/absence seizures with developmental delay (Satterstrom et al. 2020; Heyne et al. 2018). POGZ therefore regulates embryonic expression of genes that promote synapse formation and function. Furthermore, the list of up-regulated genes includes two imprinted genes *Nespas* (2.6- fold) and *Rian* (2.4-fold) (Laukoter et al. 2020; Gregg et al. 2010) which suggests a possible role for POGZ in the silencing of imprinted gene loci.

We validated RNA-seq results by ISH in E13.5 *Pogz^−/−^*, selecting genes based on significance and fold change, as well as involvement in synapse formation and function. ISH for *Pcdh11x* showed a strong reduction in gene expression in neurons of the hypothalamus, medial prefrontal cortex, and piriform cortex (Figure 2F, S4A). ISH of *Fstl5* (Follistatin-like 5), a gene involved in the Wnt/beta-catenin pathway (D. Zhang et al. 2015) whose function has not been studied in neurons, showed reduction in gene expression in both cortical neurons and progenitors in the basal ganglia (Figure 2G). *Slitrk1* and *Slitrk5* were expressed in neurons of the cortex and basal ganglia and were greatly reduced in *Pogz^−/−^* (Figure 2H, 2I, S4E, and S4F)*. Lrfn5* and *Lphn3* were expressed in cortical neurons and greatly reduced in *Pogz^−/−^* (Fig 2E, 2J, S3F, S4B). *Flrt2* and *Gabra2* were expressed in neuronal layers of cortex and the lateral ganglionic eminences, where expression was reduced in *Pogz^−/−^* (Figure 2K, 2L, S4C and S4D). Of the up-regulated genes, we validated *Slc6a1* and *Ccnd1* whose expression was increased in cortex (Figure 2D, S3D, S3E).

### POGZ promotes local chromatin accessibility at loci proximal to downregulated genes

To investigate whether POGZ regulates chromatin to modify gene expression, we compared the number and locations of open chromatin regions (OCRs) in wild type and *Pogz^−/−^* using ATAC-seq in E13.5 cortex and basal ganglia. *Pogz^−/−^* had a small reduction in OCRs (95,996 in wildtype cortex, 91,961 *Pogz^−/−^* cortex; 82,699 in wildtype basal ganglia, 76,524 in *Pogz^−/−^* basal ganglia)(Figure S5A). To assess whether *Pogz^−/−^* altered H3K27ac, a histone modification enriched at active enhancers and promoters (Rada-Iglesias et al. 2011), we performed H3K27ac ChIP-seq in wildtype and *Pogz^−/−^* cortex. Again, the mutation did not lead to overt genome-wide changes in H3K27ac peak numbers or H3K27ac levels at OCRs (Figure S5A).

On the other hand, we found that *Pogz^−/−^* have decreased chromatin accessibility at specific gene loci. Comparing normalized ATAC-seq signal in *Pogz^−/−^* versus wildtype, we observed reduction of chromatin accessibility specifically around down-regulated genes. No such changes were found around up-regulated genes. For example, across Chromosome 14, the greatest decreases in relative chromatin accessibility in *Pogz^−/−^* occurred around the down-regulated *Slitrk1,5,6* cluster (Figure 3A); on Chromosome 5, the greatest decreases were localized around the *Gabra2, Gabrg1*, and *Gabra4* gene group, as well as *Lphn3* (Figure 3C). Other examples include on Chromosome 3 and the X chromosome around the *Fstl5* and *Pcdh11x* gene loci, respectively (Figure S5D, F). Thirteen of the top fifteen down-regulated genes were found in 100 kb genomic bins that contain a 50% or greater decrease in chromatin accessibility in *Pogz^−/−^.* Notably, seven of the top fifteen down-regulated genes are found in proximity (within 100 kb) to other down-regulated genes: *Pcdh11x* and *Nap1l3, Slitrk1, 5, and 6*, and *Gabra2, Gabrg1*, and *Gabra4* are found in clusters with reduced chromatin accessibility in *Pogz^−/−^*.

**Figure 3).**
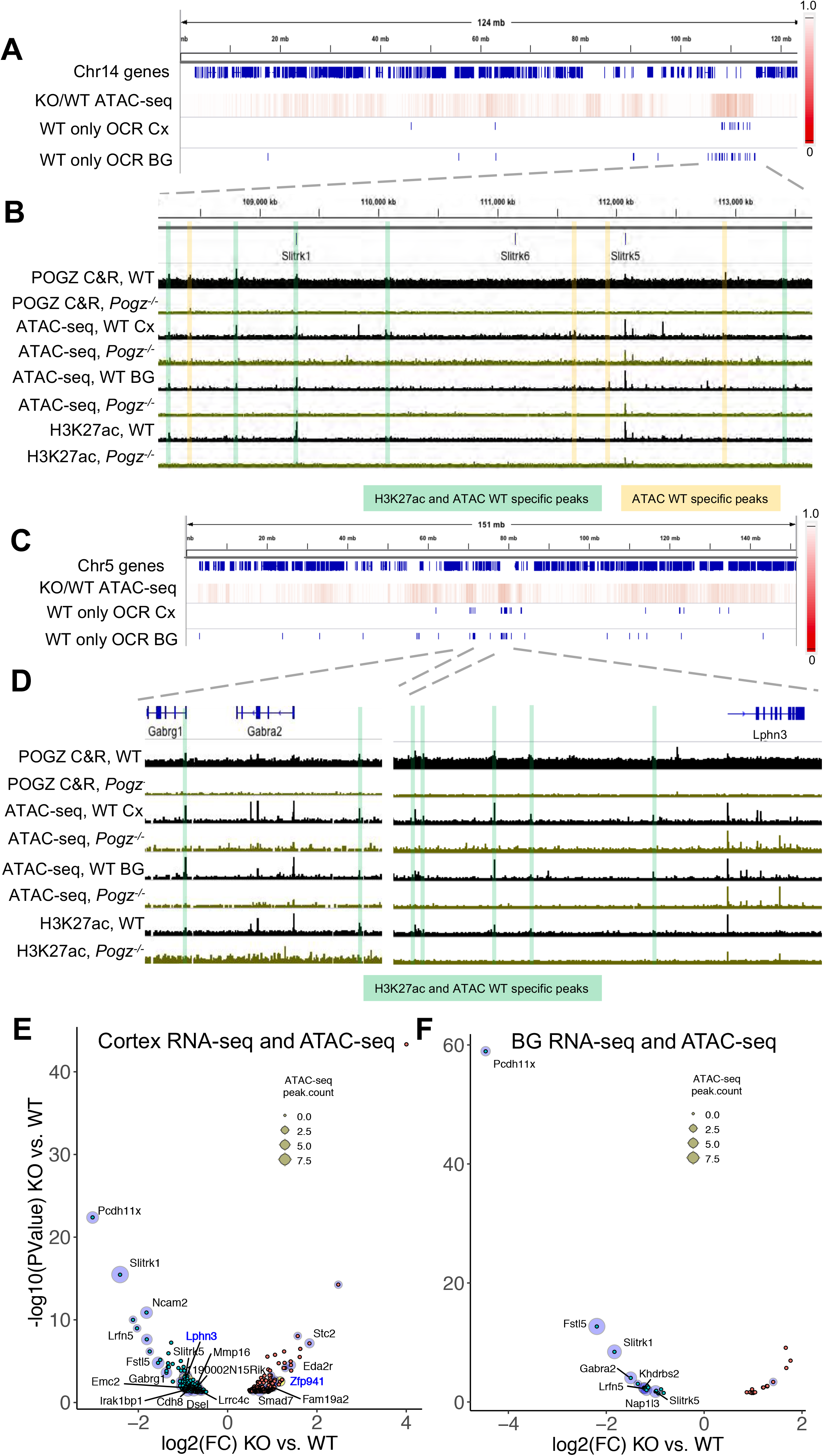
Localized changes in chromatin accessibility at DE gene loci in *Pogz^−/−^*. A) Genome browser view of all genes on mouse chromosome 14. Heatmap of ATAC-seq reads in *Pogz^−/−^* cortex normalized to wildtype in 10 kb genomic bins. Indicated below in blue hashes are peak calls of differentially accessible OCRs in wildtype cortex (Cx) and basal ganglia (BG) compared to *Pogz^−/−^*. B) Zoom in on *Slitrk1,5,6* gene cluster. Highlighted are POGZ occupied loci that are WT specific OCRs (yellow) or have both WT specific OCRs and WT specific enrichment of H3K27ac (green). Tracks are sequencing reads from individual experiments on wildtype (black) and *Pogz^−/−^* (green): Pogz C&R from E13.5 telencelphalon, ATAC-seq from E13.5 cortex (cx) and basal ganglia (bg), and H3K27ac ChIP-seq reads from E13.5 cortex. C) Genome browser view of all genes on mouse chromosome 5. Heatmap of ATAC-seq reads in *Pogz^−/−^* cortex normalized to wildtype in 10 kb genomic bins. Indicated below in blue hashes are peak calls of differentially accessible OCRs in wildtype cortex (Cx) and basal ganglia (BG) compared to *Pogz^−/−^*. D) Zoom in on *Gabrg1* and *Gabra2* gene cluster, and on *Lphn3* gene locus. Highlighted in green are POGZ occupied loci that are WT specific OCRs and have WT specific enrichment of H3K27ac. Tracks are sequencing reads from individual experiments on wildtype (black) and *Pogz^−/−^* (green): Pogz C&R from E13.5 telencephalon, ATAC-seq from E13.5 cortex (cx) and basal ganglia (bg), and H3K27ac ChIP-seq reads from E13.5 cortex. E,F) Volcano plots of RNA-seq data from differentially expressed genes at E13.5 *Pogz^−/−^* cortex (E) and basal ganglia (F). Up-regulated (red dot) or down-regulated (green dot) genes that contain proximal ATAC-seq peaks that are KO or WT specific, respectively, are indicated with blue halos. Halos indicates the number of proximal ATAC-seq peaks that are gained or lost in *Pogz^−/−^*.

The decrease in chromatin accessibility observed at down-regulated gene clusters is due to the loss of OCRs in *Pogz^−/−^.* Differential peak calling analysis identified 277 OCRs that were WT specific, and 1,835 OCRs that were *Pogz^−/−^* specific in cortex (Table S3). GO analysis on genes nearest the WT specific OCRs in cortex and basal ganglia yielded terms such as “regulation of synapse assembly” and “axon development”. (Figure S5B). Downregulated genes are proximal to WT specific OCRs, and a subset of up-regulated genes are proximal to *Pogz^−/−^* specific OCRs (Figure 3E,F).

Next, we identified 1,973 WT specific and 521 *Pogz*^−/−^ specific H3K27ac ChIP-seq peaks in cortex by differential peak calling (Table S3). GO analysis of WT specific H3K27ac peaks finds terms such as “positive regulation of synapse assembly” (Figure S5C). Indeed, at the *Slitrk1,5,6* locus we observe five WT specific H3K27ac peaks that overlap WT specific OCRs and POGZ C&R peaks (Figure 3B). Other down-regulated genes in *Pogz*^−/−^ are also proximal to POGZ-occupied loci that were WT specific OCRs: these include *Lphn3, Gabra2, Gabrg1, Fstl5, Nap1l3*, and *Pcdh11x* (Figure 3E, Figure S5E, Figure S5G). A subset of WT specific OCRs bound by POGZ were also WT specific H3K27ac peaks (Figure 3, Figure S5)

We tested POGZ C&R peaks that were WT specific OCRs and H3K27ac^+^ for enhancer activity. Two elements at the *Slitrk1,6,5* locus had enhancer activity in a luciferase transcription assay in primary cultures from embryonic cortex and basal ganglia (Figure S6A,B). Two elements proximal to the *Lphn3* locus had enhancer activity as well (Figure S6C,D). Co-transfection of POGZ did not affect candidate RE activity (data not shown), suggesting that POGZ activity at REs may depend on the local chromatin environment. These results provide evidence that these elements are enhancers whose chromatin accessibility is positively regulated by POGZ.

### POGZ and ADNP form a complex with HP1γ and co-occupy loci

Our observation that POGZ promotes chromatin accessibility and gene expression suggests that its dominant functional interaction may not be with repressive heterochromatin. Previously POGZ has been shown to interact with heterochromatin protein 1 (HP1), a structural component of heterochromatin (Nozawa et al. 2010). HP1 variants α and β localize primarily in heterochromatin, while γ is found in both heterochromatin and euchromatin and at actively transcribed genes (Canzio, Larson, and Narlikar 2014; Vakoc et al. 2005; Minc, Courvalin, and Buendia 2000). We asked which HP1 variant POGZ binds in nuclear extracts of E13.5 mouse cortex. By co-immunoprecipitation (co-IP) we found HP1α antibody did not pull down POGZ, whereas HP1γ antibody did (Figure 4A).

**Figure 4).**
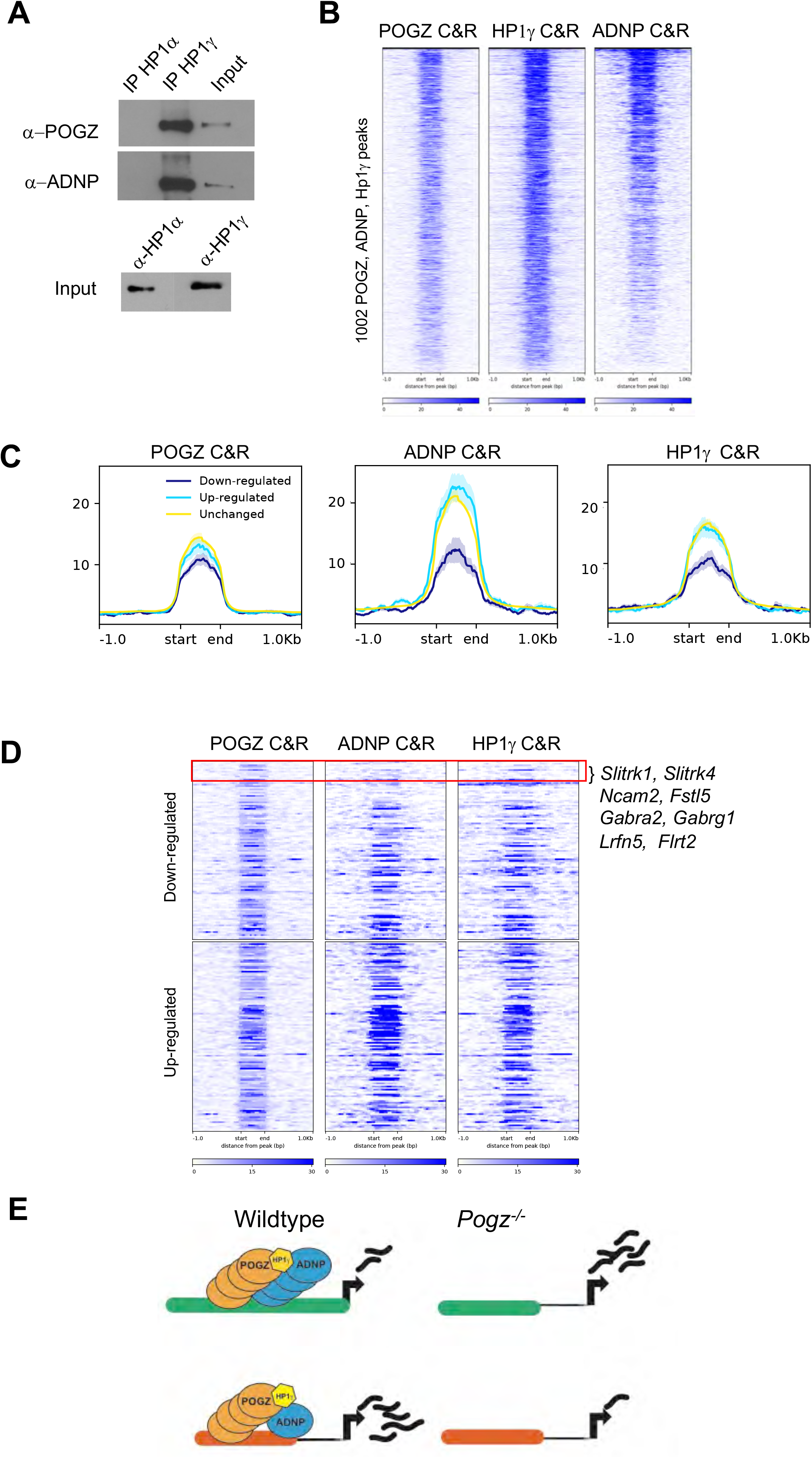
POGZ and ADNP form a nuclear complex with HP1 γ. 4A) Western blots for POGZ and ADNP in IPs for HP1α and HP1γ from nuclear extracts of E13.5 wildtype mouse cortex 4B) Heat map of POGZ, ADNP, and HP1γ C&R reads across consensus peaks for all three antibodies in E13.5 mouse forebrain. Each row represents a 2 kb window centered on peak midpoints 4C) POGZ, ADNP, and HP1γ C&R signal over POGZ C&R peaks in three different segments of the genome: proximal (within 100 kb) to down-regulated genes (qvalue<0.01), proximal to up-regulated genes (qvalue<0.01), proximal to unchanged genes. 4D) Heat map of POGZ, ADNP, and HP1γ C&R reads across consensus peaks proximal (within 100 kb) to differentially expressed genes in *Pogz^−/−^*. Each row represents a 2 kb window centered on peak midpoints, sorted by fold decrease or increase of proximal gene in *Pogz^−/−^.* 4E) Proposed model for POGZ, ADNP, and HP1γ activity at REs. High levels of binding of all three proteins leads to repression of proximal genes (observed up-regulation in *Pogz^−/−^*), while high levels of POGZ and reduced levels of ADNP and HP1γ leads to activation of proximal genes (and observed down-regulation in *Pogz^−/−^*).

HP1γ interacts with ADNP and CHD4 to form the ChAHP complex that represses gene transcription by generating inaccessible chromatin (Ostapcuk et al. 2018). ADNP is a high confidence ASD risk gene (Satterstrom et al 2019) that recognizes DNA motifs through its homeodomain and directs binding of the ChAHP complex to euchromatin. We tested whether HP1 variants interact with ADNP in E13.5 cortex. As we saw for POGZ, HP1α did not co-IP with ADNP, whereas HP1γ did (Figure 4A).

Next, we assessed whether POGZ, HP1γ and ADNP occupy the same chromosomal loci in E13.5 mouse telencephalon using the C&R assay with anti-HP1γ and anti-ADNP antibodies. Computational analysis found 1,002 consensus loci occupied by all three proteins across duplicate experiments (Figure 4B, S7A). ATAC-seq and ChIP-seq from E13.5 cortex showed these loci are euchromatic, with accessible chromatin and enriched for H3K27ac, and had no detectable H3K9me3 heterochromatin (S7B). These loci were located at transcription start sites and enriched for GO terms such as “pallium development”, “gliogenesis”, and “ephrin receptor signaling”, which suggests that shared targets are proximal euchromatic elements of genes that regulate cortical neurodevelopment (Figure S7C, S7D).

As ADNP is a repressor of neuronal gene expression (Ostapcuk et al. 2018), we asked whether ADNP is present where POGZ promotes neuronal gene expression by RNA-seq. Interestingly, there is reduced co-occupancy of ADNP and HP1γ with POGZ at loci proximal to genes down-regulated in *Pogz^−/−^*, compared to genes that were up-regulated or unchanged (Figure 4C, 4D). Examples of POGZ occupied loci where ADNP and HP1γ binding was reduced are the *Slitrk* and *Gabra2* genes (Figure 4D). We propose a model wherein POGZ acts as a positive regulator of transcription when it occupies loci with reduced co-occupancy of ADNP and HP1γ, but acts as a repressor with high co-occupancy of ADNP and HP1γ (Figure 4E).

### *Pogz* heterozygote mice have reduced *Adnp* expression

Heterozygous loss-of-function mutations in *POGZ* lead to ASD, ID and other neurodevelopmental phenotypes in affected individuals. Consequently we were interested in whether we could detect a phenotype in *Pogz^+/−^* mice that express half the amount of POGZ protein (Figure S8A,B). To look for gross anatomical defects, we performed MRI on P28 *Pogz^+/−^* brains and found no significant change in cortex size (Figure S8C).

*Pogz^+/−^* mice have been described to have electrophysiological phenotypes in mPFC (Cunniff et al. 2020); thus, we analyzed mPFC of P28 *Pogz^+/−^*. We performed RNA-seq in *Pogz^+/−^* and wildtype mice, and found 29 significantly (qvalue<0.05) down-regulated and ten up-regulated genes across three replicates per genotype (Table S4). *Adnp* was one of the most highly down-regulated genes (1.5-fold, q=0.0008)(Figure 5A,B). ISH experiments confirmed reduction of *Adnp* mRNA levels in *Pogz^+/−^* cortex (Figure 5C).

**Figure 5).**
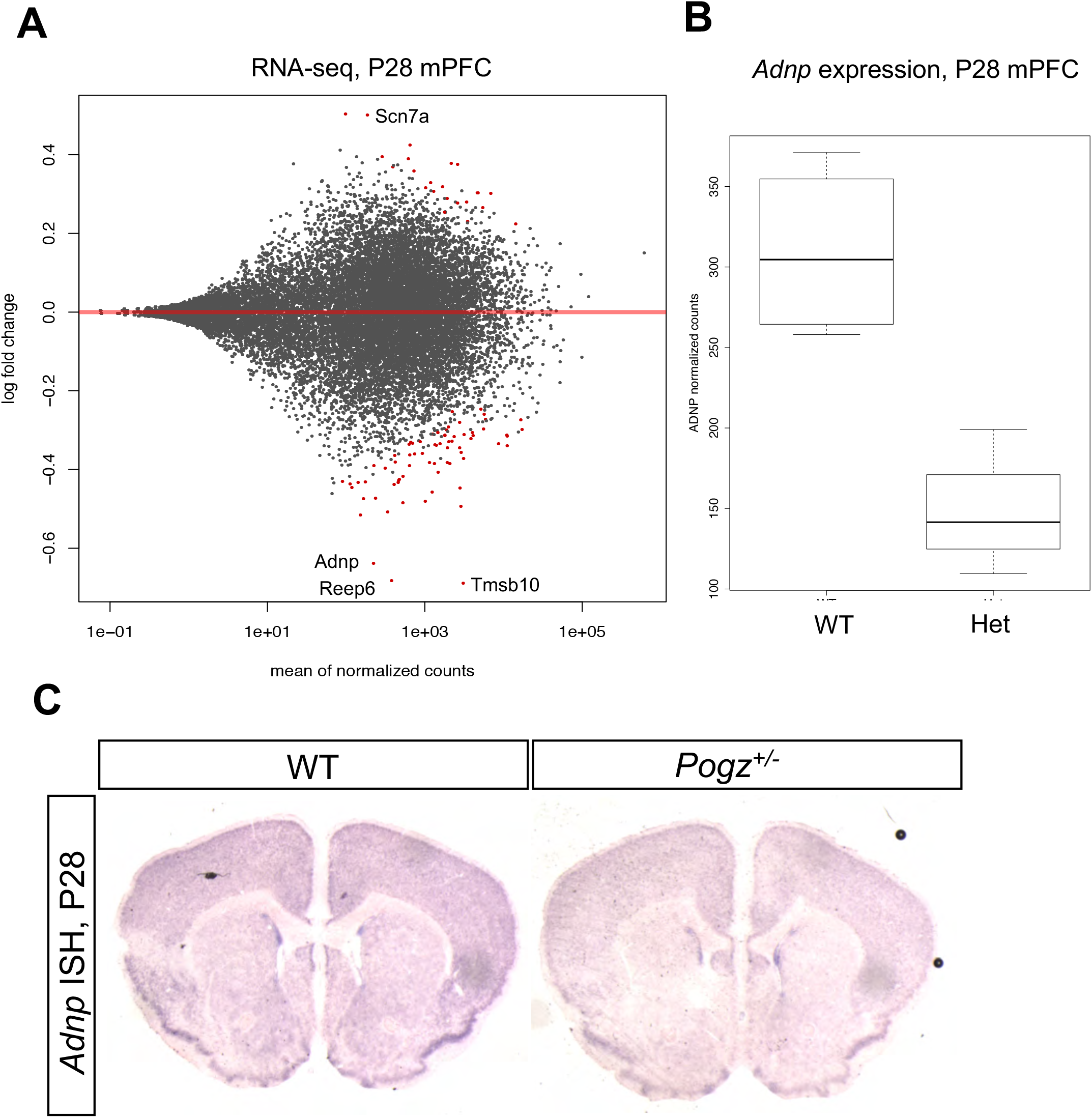
*Adnp* expression reduced in *Pogz* heterozygote. 5A) MA plot showing differentially expressed genes in P28 *Pogz^+/−^* mPFC RNA-seq counts compared to wildtype. Significantly (qvalue<0.05) differentially expressed genes are indicated in red dots. 5B) Normalized *Adnp* RNA-seq read counts in wildtype and *Pogz^+/−^* mPFC. 5C) ISH using antisense *Adnp* RNA probe in wildtype and *Pogz^+/−^* P28 coronal sections.

### POGZ-occupied loci in human fetal cortex are euchromatic and enriched for NDD genes

*POGZ* is expressed in the mid-fetal human cortex (Figure 6A). To test whether POGZ binds HP1α or HP1γ we performed co-IP on nuclear extracts from 17 gestation week (gw) cortex. As in mouse, HP1α antibody did not pull down POGZ, whereas HP1γ antibody did (Figure 6B). We studied POGZ occupancy in nuclei isolated from 17 gw and 18 gw human cortex and identified 9,089 consensus POGZ C&R peaks across three replicates. POGZ-occupied loci exhibited high levels of chromatin accessibility in data from published human fetal cortex ATAC-seq (Markenscoff-Papadimitriou et al. 2020). As in the mouse, C&R for HP1γ and ChIP-seq for histone modifications show POGZ and HP1γ primarily overlapped H3K27ac peaks and not H4K20me3 heterochromatin modifications (Figure 6C). Paralleling findings in mouse cortex, POGZ was localized at distal loci in the *SLITRK1,5,6* cluster (data not shown). These experiments provide evidence that POGZ co-occupies euchromatic and heterochromatic loci with HP1γ in both human and mouse.

**Figure 6).**
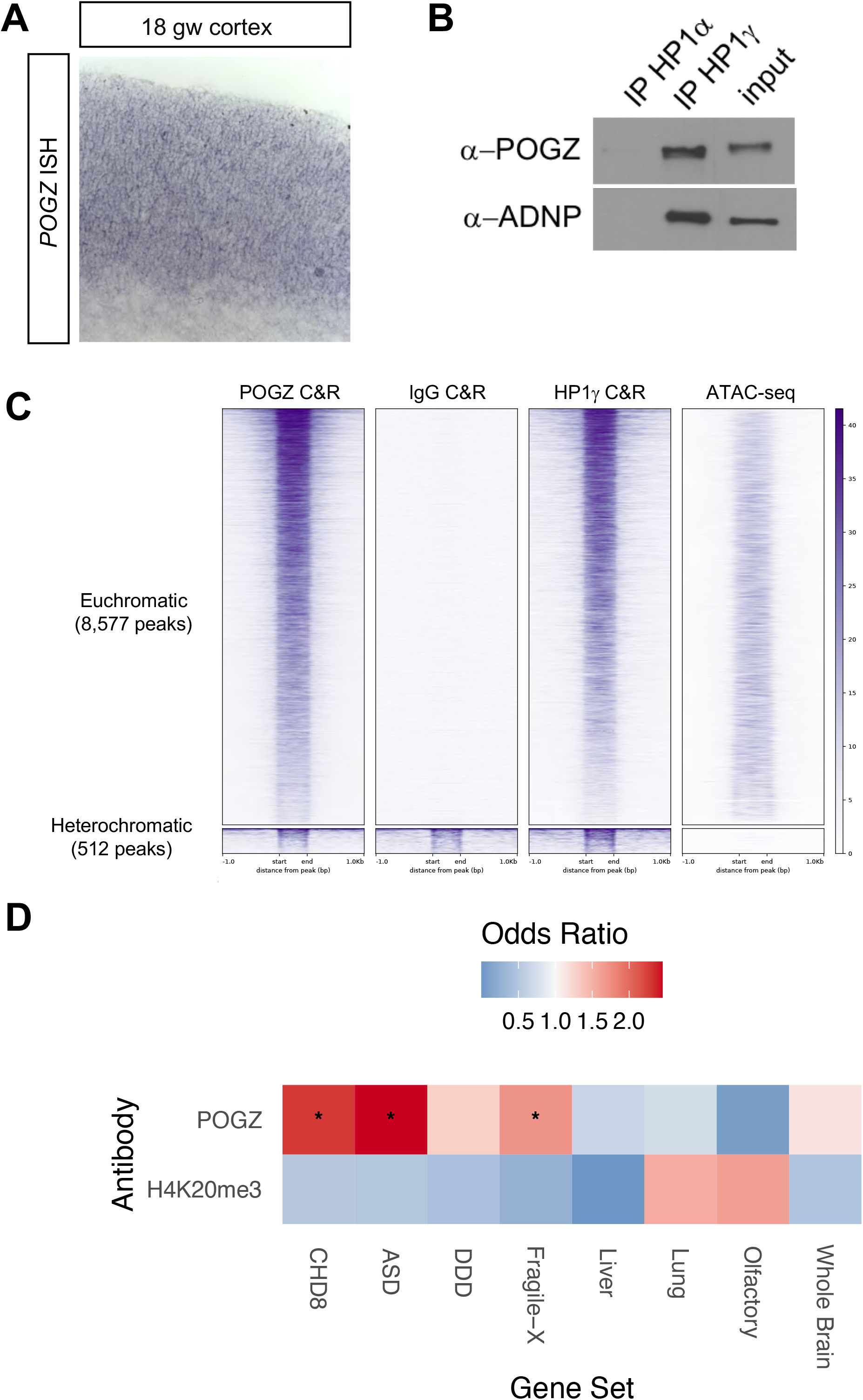
POGZ binds euchromatic loci and HP1γ in human fetal cortex. 6A) ISH of *POGZ* antisense probe (purple) in 18 gw fetal cortex tissue 6B) Co-IPs for HP1α and HP1γ from nuclear extracts of 18 gw cortex. 6C) Heat map of POGZ, HP1γ, and IgG control C&R reads across POGZ consensus peaks (*n* = 9,089) in human mid-fetal cortex. Each row represents a 2 kb window centered on peak midpoints, sorted by the POGZ C&R signal. ATAC-seq from mid-fetal cortex sampls shown. Euchromatic peaks on the top row, heterochromatic peaks on the bottom row. RPM, reads per million. 6D) Gene set enrichment analysis for genes proximal (within 50 kb) to POGZ C&R peaks, H4K20me3 peaks as control. Gene sets that are significantly enriched in POGZ C&R-proximal genes are indicated with a star (qvalue <0.001 after multiple testing correction). Gene sets included ASD risk genes (Satterstrom et al. 2020), biological targets of fragile X mental retardation protein (FMRP)(Darnell et al. 2011), biological targets of ASD gene CHD8 (Cotney et al. 2015), and developmental delay disorder (DD) genes (Deciphering Developmental Disorders Study 2017). See STAR Methods for description of control gene sets.

**Figure 7).**
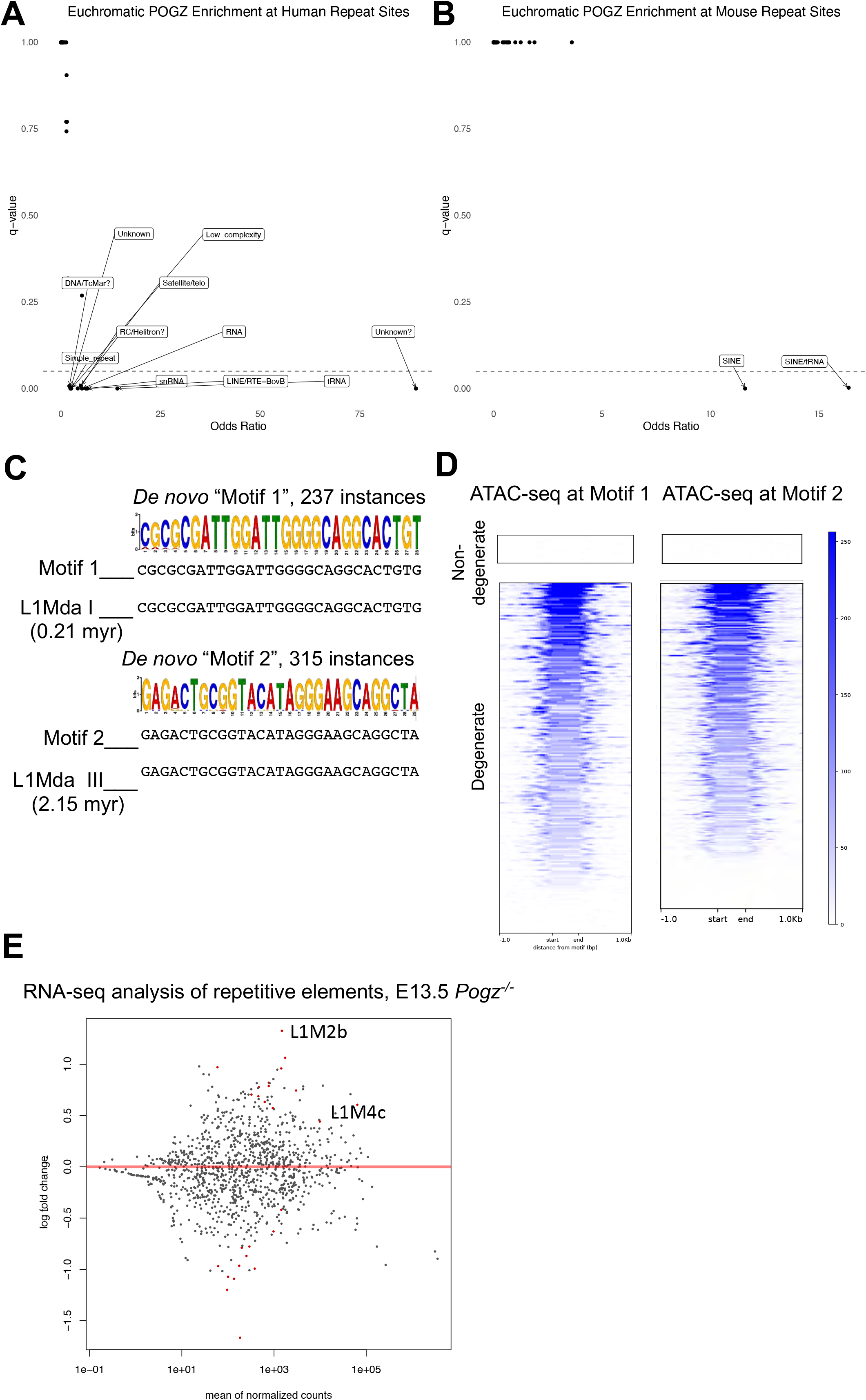
POGZ occupied loci enriched for TEs, L1 retrotransposon motifs. 7A,B) Volcano plots of transposable elements enriched in euchromatic POGZ C&R peaks from human 18 gw cortical (A) and mouse E13.5 telencephalon (B) samples. Elements under the dotted line (qvalue <0.05 after multiple testing correction) are significantly enriched. 7C) Repeatmasker alignment of two *de novo* POGZ binding motifs identified by MEME-CHIP analysis of POGZ C&R data. 7D) Plot and Heatmap of ATAC-seq signal at instances of predicted POGZ DNA-binding motifs 1 and 2 in POGZ C&R peaks. POGZ peaks are clustered by whether there is no mismatch with the motif sequence (non-degenerate) or whether there are mismatches (degenerate). 7E) MA plot showing differentially expressed repetitive elements in *Pogz^−/−^* E13.5 RNA-seq data, significantly (qvalue<0.05) differentially expressed genes are indicated in red dots.

Next, we asked whether POGZ occupies gene loci implicated in NDDs. We performed gene set enrichment analysis for genes proximal (within 50 kb) to POGZ C&R peaks in human mid-fetal cortex. We tested various NDD relevant gene sets: ASD and Developmental Delay risk genes (Satterstrom et al. 2020; Deciphering Developmental Disorders Study 2017), and targets of the ASD gene CHD8 and Fragile-X Syndrome gene FMRP (Cotney et al. 2015; Darnell et al. 2011), and compared enrichment to control gene sets. POGZ C&R peak-proximal genes were significantly enriched for ASD risk genes and targets of CHD8 and FMRP (qvalue<0.001, Benjamini-Hochberg multiple test correction), while they were not enriched for whole brain expressed genes (q value = 0.756). Furthermore, the proportion of CHD8 target genes proximal to POGZ C&R peaks was statistically significant compared to the proportion of 1,000 random samples of whole brain expressed genes (empirical p-value = 0; see Methods). From these analyses we concluded that POGZ occupied loci in the mid-fetal human cortex are significantly enriched for ASD risk genes and CHD8 targets.

### POGZ occupied loci are enriched for Transposable Elements (TEs)

Because POGZ contains a *pogo* transposon sequence in the final exon, we wondered whether POGZ occupied loci are enriched for TEs. We asked if any classes of TEs are enriched in euchromatic POGZ C&R loci from the developing mouse telencephalon and human cortex. In mouse, SINE elements and tRNA-derived SINEs were enriched (qvalue<0.05 after multiple testing correction); in human, LINE elements and an unknown repetitive element were enriched (qvalue<0.05) (Figure 7A,7B).

*De novo* motif analysis of mouse POGZ C&R peaks identified one 28bp and one 29bp motif that mapped to the promoters of L1Mda-1 and L1Mda-III, two of the most recently evolved L1 LINE genes in the mouse genome (Figure 7C) (Sookdeo et al. 2013). Non-degenerate instances of these motifs in the C&R peaks are heterochromatic, while degenerate instances of these motifs are euchromatic and have accessible chromatin, suggesting they are at REs (Figure 7D).

Given the enrichment of L1 motifs in POGZ bound sites, we wondered whether POGZ regulates L1 transcription. We revisited *Pogz^−/−^* RNA-seq data from E13.5 cortex and analyzed differences in transcription of repetitive elements. Eight genes had significantly increased expression (qvalue<0.05) in *Pogz^−/−^*, including two L1 genes; the most up-regulated gene (2.5 fold) was the retrotransposon L1M2b (Figure 7E). Taken together, our analyses conclude that POGZ C&R loci are enriched for TEs and L1 sequence motifs, and that POGZ modulates L1 transcription in developing mouse cortex.

## Discussion

Mutations in POGZ are strongly associated with risk for NDDs such as ASD, ID, and Developmental Delay (Satterstrom et al. 2020; Deciphering Developmental Disorders Study 2015). Here, we generated a constitutive *Pogz* null allele to study POGZ’s molecular function in mouse neuronal development. We performed chromatin and gene expression profiling in *Pogz^−/−^* embryos, and mapped POGZ occupancy genome-wide in embryonic mouse and fetal human tissue. Our analyses conclude POGZ binding to promoters and distal enhancers promotes chromatin accessibility and transcription at a specific set of genes, many of which encode synaptic proteins. We provide evidence that this function is distinct from when POGZ is found in a nuclear complex with HP1γ and ADNP, another high confidence ASD risk gene and gene repressor (Ostapcuk et al. 2018). Below, we discuss the implications of our findings on POGZ function and speculate on an evolutionary role of POGZ.

### POGZ regulates euchromatin

POGZ is frequently cited as a heterochromatin-associated protein and a transcriptional repressor. One of the main conclusions of our analysis is that POGZ occupies euchromatic as well as heterochromatic loci and acts to promote as well as repress transcription. Genome-wide mapping of POGZ occupancy by C&R in developing mouse (Figure 1) and human (Figure 6) brain tissues revealed it predominantly binds euchromatic loci. POGZ C&R peaks are found at transcription start sites and distal REs with accessible chromatin and the active RE mark H3K27ac. Luciferase assays validated the transcription enhancer function of two distal REs bound by POGZ; furthermore, 96 Vista enhancers with activity in embryonic brain tissue overlap POGZ C&R peaks. Thus, like the ASD risk genes CHD8 and ADNP (Cotney et al. 2015; Ostapcuk et al. 2018), POGZ binds REs.

POGZ’s reputation as a transcriptional repressor is due to a report that it binds HP1α, a chromatin scaffolding protein associated with repressive heterochromatin (Nozawa et al. 2010). We found POGZ preferentially binds HP1γ in co-IP experiments we performed using nuclear extracts from developing mouse and human brain (Figure 4, Figure 6). HP1γ is a variant of HP1 associated with both heterochromatin and euchromatin. Missense *de novo* ASD-associated variants identified in the HP1-binding zinc finger domain of POGZ (Sanders et al. 2015; Satterstrom et al. 2020; Stessman et al. 2016) may point to POGZ’s interaction with HP1γ being relevant to the pathology of ASD. Future work should focus on studying deleterious human mutations to identify which domains of POGZ are directly relevant to the ASD phenotype.

### POGZ promotes neuronal gene transcription and chromatin accessibility at gene clusters

One of the more notable findings of our study was that POGZ promotes the expression of genes involved in synapse formation and function. RNA-seq and ISH analysis of *Pogz^−/−^* identified genes encoding GABA_A_ receptor subunits, LPHN3/FLRT synapse complex, and leucine-repeat rich proteins as highly down-regulated (Figure 2). Strikingly, *Slitrk1,2,4* and *5* were down-regulated in cortex and basal ganglia. *Slitrk1* is a Tourette’s candidate gene (Abelson et al. 2005; O’Roak et al. 2010), and *Slitrk5* mouse mutants have OCD-like behaviors (Shmelkov et al. 2010). We identified candidate REs regulated by POGZ at these gene loci (and validated two in the *Slitrk1,6,5* locus).

Mechanistic insight into how POGZ functions to promote gene expression was gained by ATAC-seq and ChIP-seq analysis of *Pogz^−/−^* brain. Remarkably, the effects of POGZ deletion on chromatin accessibility and H3K27ac deposition were highly restricted to the gene loci that were down-regulated in *Pogz^−/−^* (Figure 3). This highly localized effect of mutation of an NDD-linked transcriptional regulator is unusual: *Arid1b, Chd8*, *Mecp2*, and *Tbr1* mutants have more diffuse effects on gene expression (Gompers et al. 2017; Fazel Darbandi et al. 2018).

A striking feature of genes whose expression, and chromatin accessibility is regulated by POGZ is that they are located frequently in gene clusters. *Slitrk1,6,5; Pcdh11x* and *Nap1l3; Gabra2* and *Gabrg1* are some examples. The observation that genes down-regulated in *Pogz^−/−^* are located in clusters, and that chromatin accessibility losses are restricted to these clusters, is reminiscent of regulation of the β-globin locus in erythroid cells. There, the locus control region (LCR) coordinates multiple β-globin genes by recruiting chromatin-modifying, coactivator, and transcription factor complexes (Fang et al. 2003). Intriguingly, POGZ is required for fetal hemoglobin expression, although direct regulation of the β-globin locus or LCR by POGZ has not been reported (Gudmundsdottir et al. 2018). We speculate that POGZ may regulate the clustered neuronal genes in a similar manner. OCRs that are regulated by POGZ in these gene clusters may be LCRs coordinating gene expression. Some of these candidate REs regulated by POGZ can drive transcription (Figure S6).

### Convergent genomic targets of ASD risk genes POGZ, ADNP, and CHD8

C&R experiments showed that half of POGZ occupied loci are co-occupied by ChAHP complex members Hp1γ and ADNP (Figure 4). ADNP is a high confidence ASD risk gene and a repressor of neuronal lineage gene expression; *ADNP*^−/−^ ES cells and mouse embryos have defects in neuronal fate specification (Ostapcuk et al. 2018; Pinhasov et al. 2003). Because ChAHP binding is markedly reduced at loci where POGZ promotes gene expression (Figure 4), we proposed a model wherein at loci that are co-occupied by POGZ and ADNP/ChAHP, ADNP antagonizes POGZ’s activating potential.

POGZ and ADNP are similar proteins, both containing zinc finger domains that bind HP1 proteins, as well as DNA binding domains (ADNP has a homeobox, POGZ has a DDE transposase domain) (Ostapcuk et al). ADNP syndrome, also known as Helsmoortel-Van Der Aa syndrome, and POGZ syndrome, also known as White-Sutton syndrome, have similar hallmark features: ID with ASD in a subset of patients, developmental delay, and characteristic craniofacial features (Batzir et al. 2020; Van Dijck et al. 2019). The fact that we find these proteins share >1,000 genomic targets in the developing telencephalon may lead us to the underlying genes, cell types, and pathways impacted in children with mutations in POGZ and ADNP.

Another point of convergence of POGZ with ASD risk genes is the enrichment of ASD genes and CHD8 targets in POGZ genomic targets in human mid-fetal cortex (Figure 6). This finding suggests that POGZ may regulate other ASD risk genes, as has been found for CHD8 (Cotney et al. 2015). Indeed, the enrichment of CHD8 occupied genes at POGZ-occupied loci suggests these two ASD risk genes and chromatin remodelers share common genomic targets. Defining the transcription networks co-regulated by POGZ, ADNP, and CHD8 will open the door to elucidating convergent mechanisms that predispose to ASD and DD.

### Transposable elements at POGZ occupied loci

One of the surprising findings of our data is that POGZ C&R bound loci in human and mouse are enriched for SINE and LINE TEs (Figure 7). Like POGZ, ADNP-occupied loci are also enriched for SINE TEs (Kaaij et al. 2019). The 28 and 29bp *de novo* motifs found in POGZ occupied loci are identical to L1 retrotransposon promoter sequences (Sookdeo et al. 2013). The presence of degenerate versions of these TE sequences in euchromatic POGZ C&R peaks suggests that these TEs may have evolved to become neurodevelopmental REs. This finding also suggests that POGZ may be regulating rates of somatic mutation. Rates of somatic mutation in neurons can be altered in NDDs; Rett syndrome models show increased L1 retrotransposition in MECP2 knockout cells (Muotri et al. 2010; D’Gama and Walsh 2018). We do not know whether POGZ regulates L1 retrotransposition, although RNA-seq analysis shows increased RNA levels of L1 LINE genes in *Pogz^−/−^*.

The POGZ gene itself contains a *pogo* family transposon sequence and is a rare example of a domesticated transposon present in vertebrates. POGZ is a chimeric gene that contains a transposase domain in the C terminus and ZNF domains in the N terminus; it evolved from a *Passer* transposon inserting into a ZNF gene (Gao et al. 2020). Interestingly, the transposase domain has retained its catalytic DD35D triad, and multiple *de novo* mutations in ASD patients have been identified in the transposase domain as well (Sanders et al. 2015; Satterstrom et al. 2020). The possibility that POGZ’s transposase domain has retained its ability to catalyze transposition events during development is an intriguing possibility that remains to be explored.

## SUPPLEMENTARY FIGURE LEGENDS

**Figure S1.**
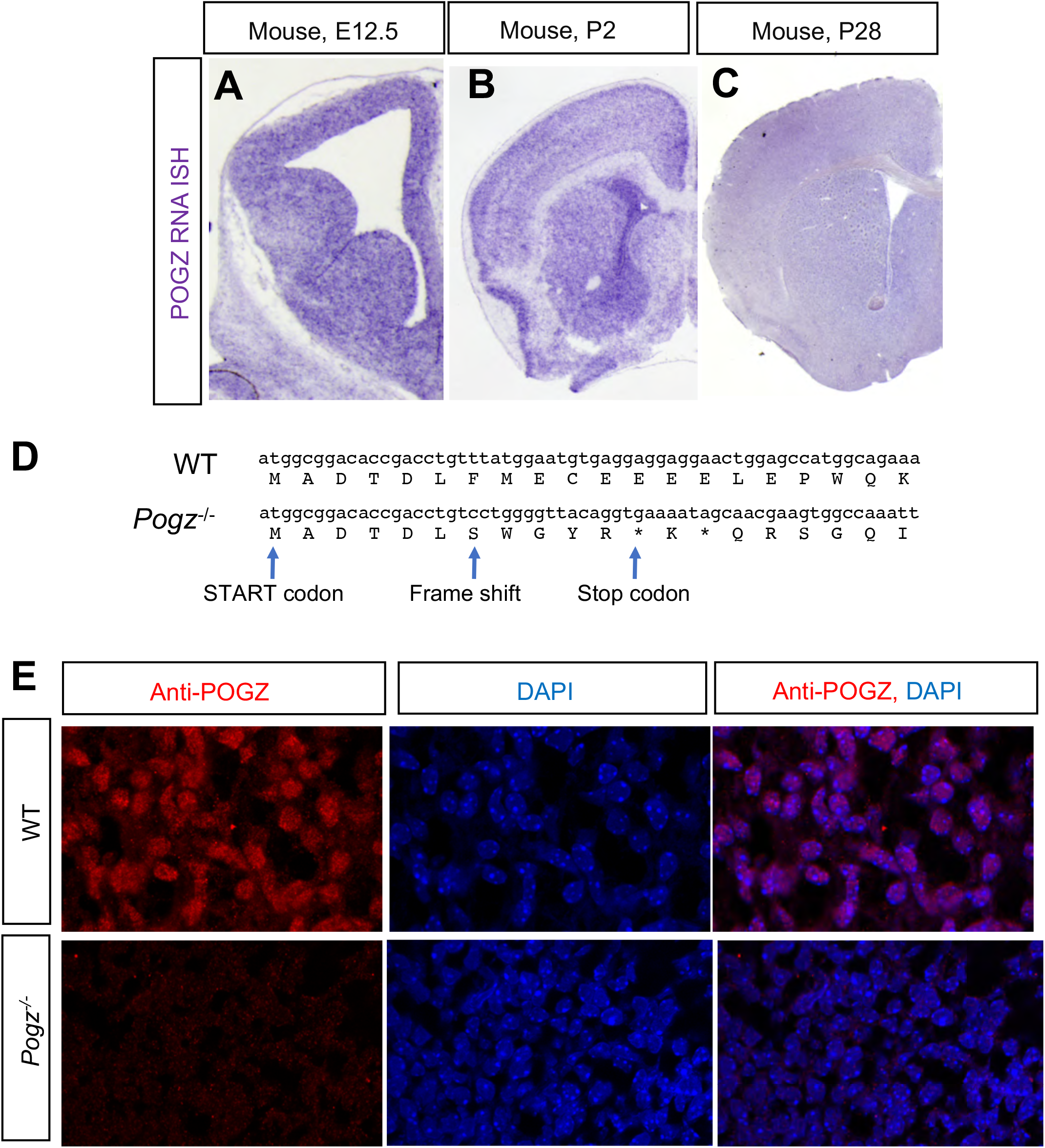
**S1A-C)** ISH with antisense *Pogz* probe of wildtype E12.5 (A), postnatal day 2 (B), and postnatal day 28 mouse forebrain (C). S1D) Sanger sequencing of wildtype and *Pogz^−/−^* alleles. Premature stop codon is indicated in *Pogz^−/−^* allele. S1E) Anti-POGZ immunostaining (red) and DAPI (blue) in E13.5 WT and *Pogz^−/−^* basal ganglia at 63x magnification.

**Figure S2.**
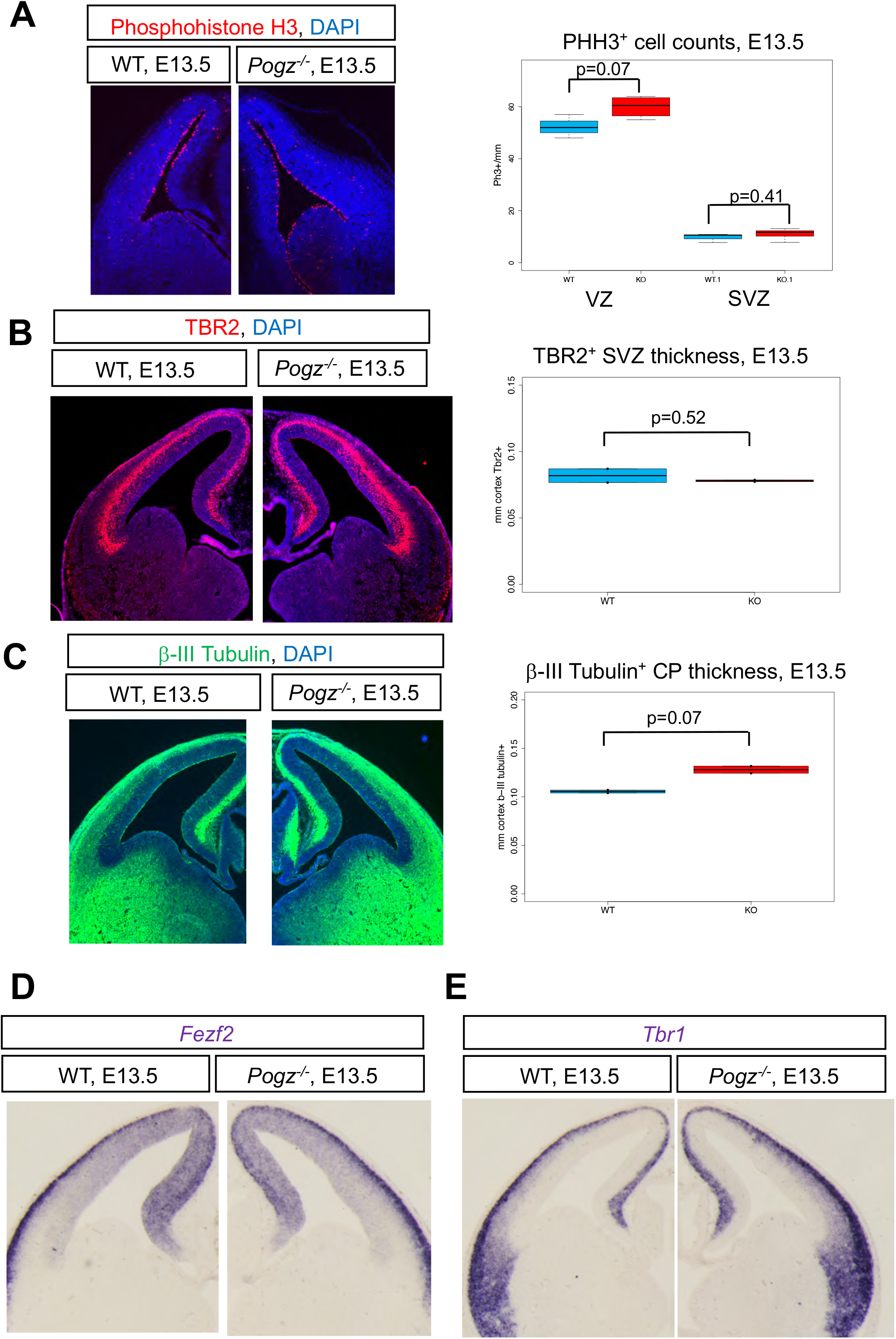
**S2A)** Anti-phosphohistone H3 immunofluorescence (Red) and DAPI (Blue) in E13.5 wildtype and *Pogz^−/−^*, and quantification of positive cells in the cortex ventricular zone (VZ) and subventricular zone (SVZ). S2B) Anti-TBR2 immunofluorescence (Red) and DAPI (Blue) in E13.5 wildtype and *Pogz^−/−^*, and quantification of SVZ layer thickness. S2C) Anti-β-III Tubulin (Tuj1) immunofluorescence (Green) and DAPI (Blue) in E13.5 wildtype and *Pogz ^−/−^*, and quantification of cortical plate (CP) thickness. S2D) Layer 5 neuron marker gene *Fezf2* ISH (purple) of E13.5 wildtype and *Pogz^−/−^*. S2E) Layer 6 neuron marker gene *Tbr1* ISH (purple) of E13.5 wildtype and *Pogz^−/−^*.

**Figure S3.**
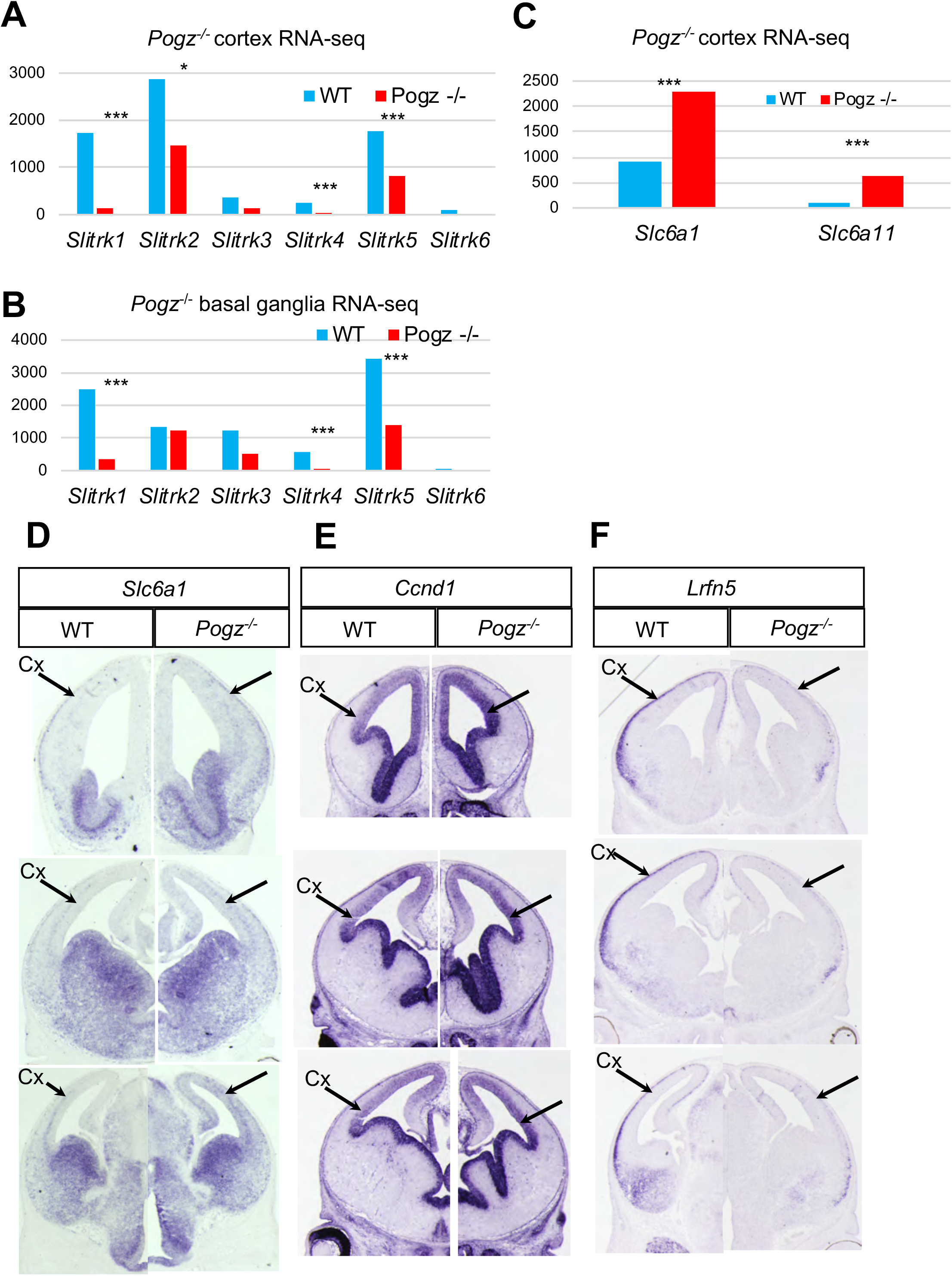
**S3A-C)** Normalized reads across RNA-seq replicates for WT and *Pogz*^−/−^ at E13.5 **S3D-F)** Coronal sections from rostral (top) to caudal (bottom) of E13.5 wildtype and *Pogz^−/−^.* ISH of genes up-regulated in *Pogz^−/−^ at* E13.5: *Slc6a1* (D), *Ccnd1*(E), and down-regulated gene *Lrfn5*(F).

**Figure S4.**
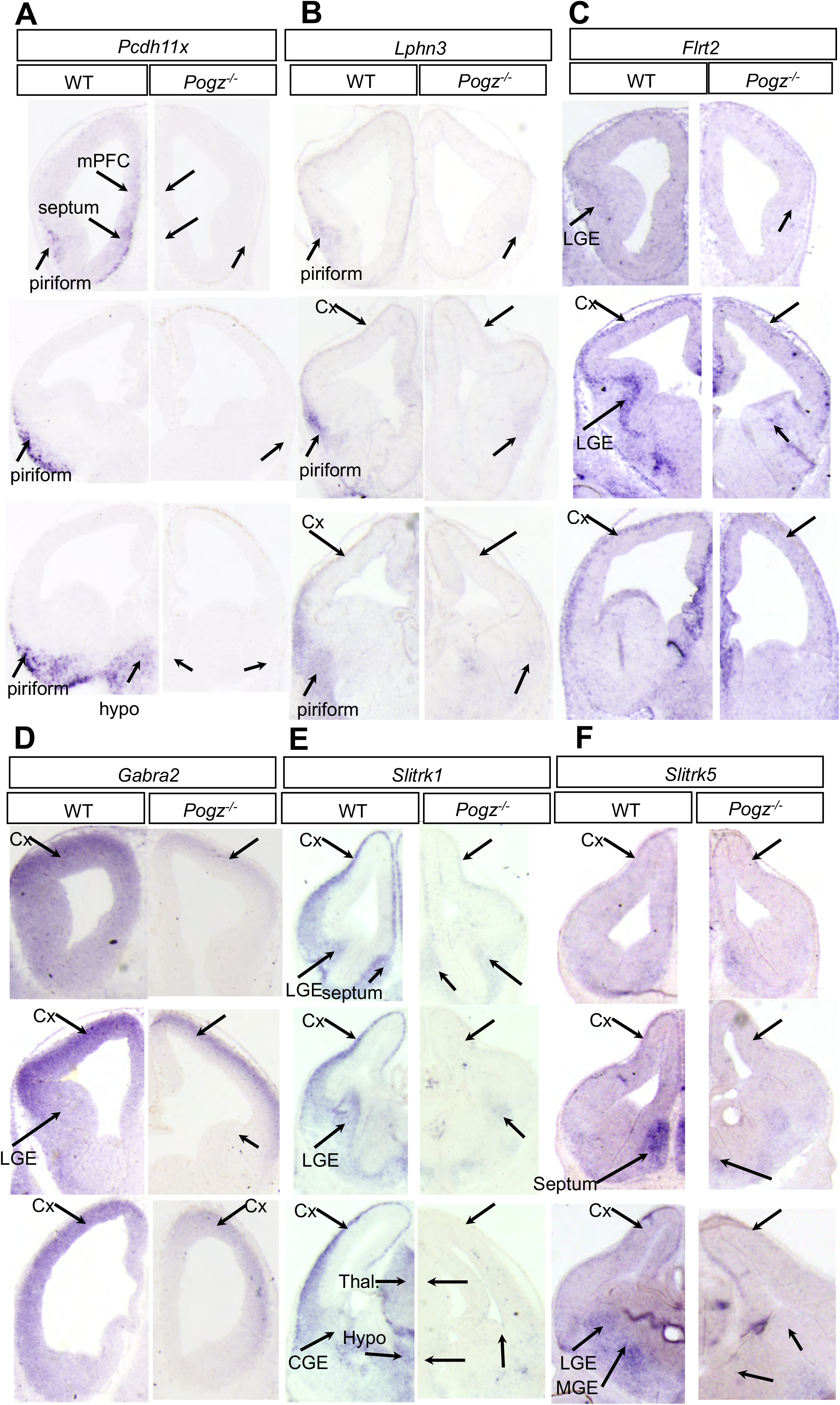
**S4A-F)** Coronal sections from rostral (top) to caudal (bottom) of E13.5 wildtype and *Pogz^−/−^* telencephalon. ISH of genes downregulated in *Pogz^−/−^ at* E13.5: *Pcdh11x* (A), *Lphn3* (B), *Flrt2* (C), *Gabra2* (D), *Slitrk1* (E), *Slitrk5* (F).

**Figure S5.**
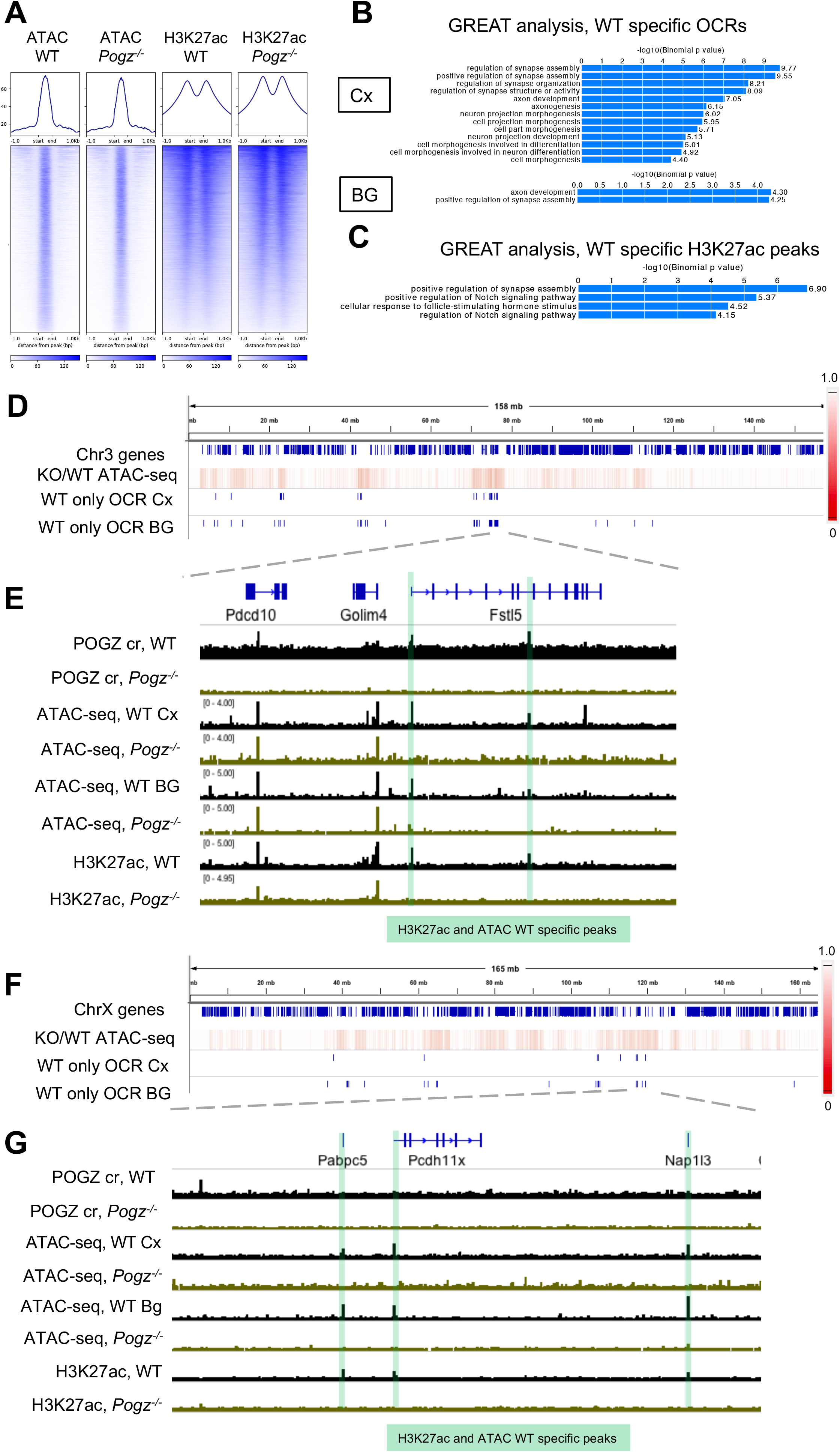
**S5A)** Plots and Heatmaps of ATAC-seq and H3K27ac signal in individual wildtype and *Pogz^−/−^* experiments at all OCRs identified in WT cortex at E13.5. S5B) GO terms significantly enriched in genes nearest to WT specific ATAC-seq peaks in E13.5 Cortex (Cx) and Basal ganglia (BG). S5C) GO terms significantly enriched in genes nearest to WT specific H3K27ac ChIP-seq peaks in E13.5 Cortex. S5D) Genome browser view of all genes on mouse chromosome 3. Heatmap of ATAC-seq reads in *Pogz^−/−^* cortex normalized to wildtype in 10 kb genomic bins. Indicated below in blue hashes are peak calls of differentially accessible OCRs in wildtype cortex (Cx) and basal ganglia (BG) compared to *Pogz^−/−^.* S5E) Zoom in on *Fstl5* gene locus. Highlighted in green are POGZ occupied loci that have both WT specific OCRs and H3K27ac enrichment. Tracks are sequencing reads from individual experiments on wildtype (black) and *Pogz^−/−^* (green): Pogz C&R from E13.5 telencephalon, ATAC-seq from E13.5 cortex (cx) and basal ganglia (bg), and H3K27ac ChIP-seq reads from E13.5 cortex. S5F) Genome browser view of all genes on mouse X chromosome. Heatmap of ATAC-seq reads in *Pogz^−/−^* cortex normalized to wildtype in 10 kb genomic bins. Indicated below in blue hashes are peak calls of differentially accessible OCRs in wildtype cortex (Cx) and basal ganglia (BG) compared to *Pogz^−/−^.* S5G) Zoom in on *Pcdh11x* and *Nap1l3* gene locus. Highlighted in green are POGZ occupied loci that have both WT specific OCRs and H3K27ac peaks. Tracks are sequencing reads from individual experiments on wildtype (black) and *Pogz^−/−^* (green): Pogz C&R from E13.5 telencelphalon, ATAC-seq from E13.5 cortex (cx) and basal ganglia (bg), and H3K27ac ChIP-seq reads from E13.5 cortex.

**Figure S6.**
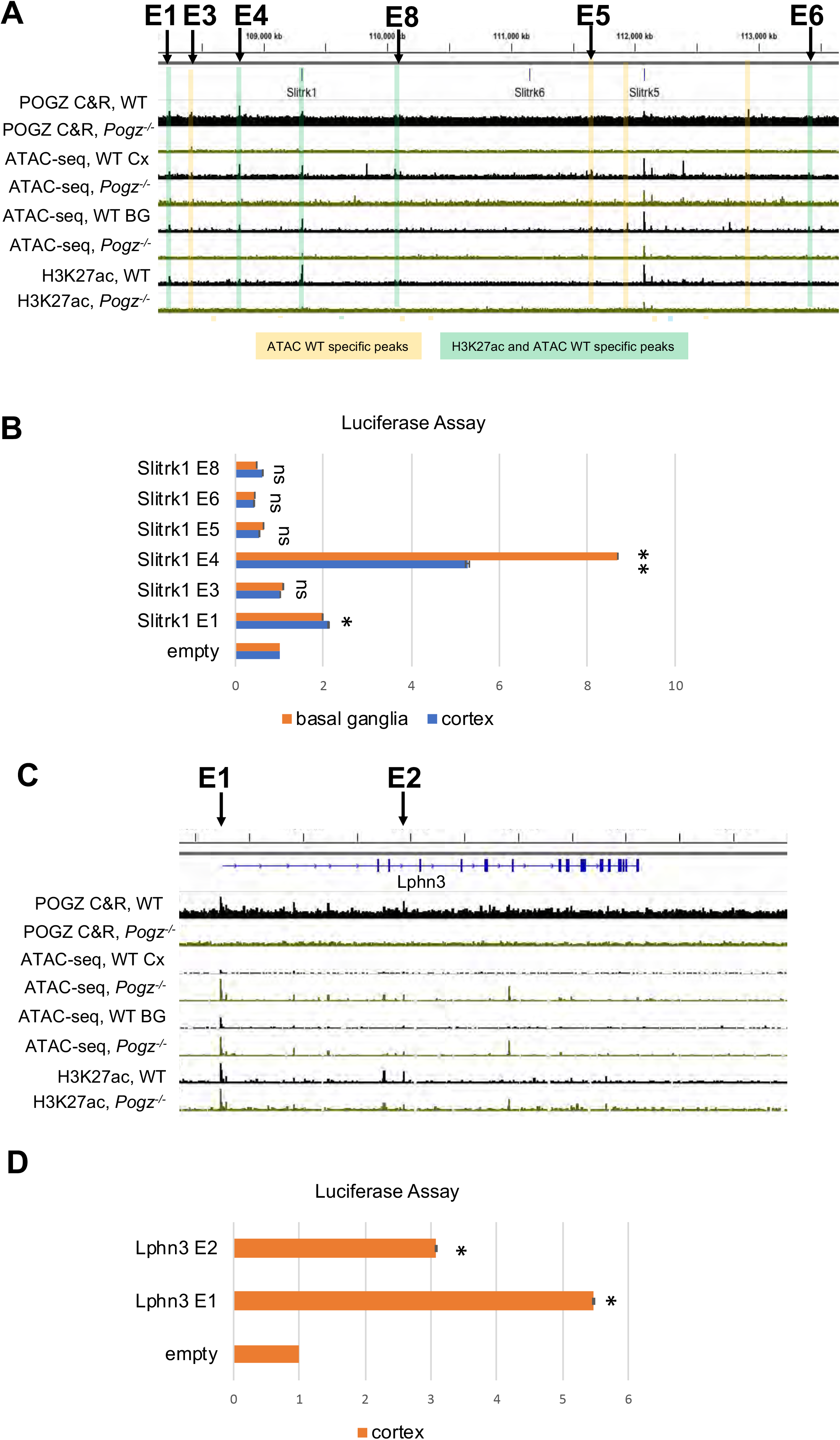
**S6A)** POGZ occupied loci in the *Slitrk1,6,5* locus that were tested by luciferase assay (indicated with arrows). Highlighted are POGZ occupied loci that are WT specific OCRs (yellow), or also have WT specific H3K27ac enrichment (green). Tracks are sequencing reads from individual experiments on wildtype (black) and *Pogz^−/−^* (green): Pogz C&R from E13.5 telencelphalon, ATAC-seq from E13.5 cortex (cx) and basal ganglia (bg), and H3K27ac ChIP-seq reads from E13.5 cortex. Experiments and peak calling were performed in duplicate or triplicate, but tracks from single experiments are shown. S6B) Mean firefly luciferase levels in primary cortical and basal ganglia cultures normalized to Renilla, normalized to empty vector, for each enhancer candidate in *Slitrk1,6,5* locus. S6C) POGZ occupied loci in the *Lphn3* locus that were tested by luciferase assay (indicated with arrows). S6D) Mean firefly luciferase levels in primary cortical cultures normalized to Renilla, normalized to empty vector, for each enhancer candidate.

**Figure S7.**
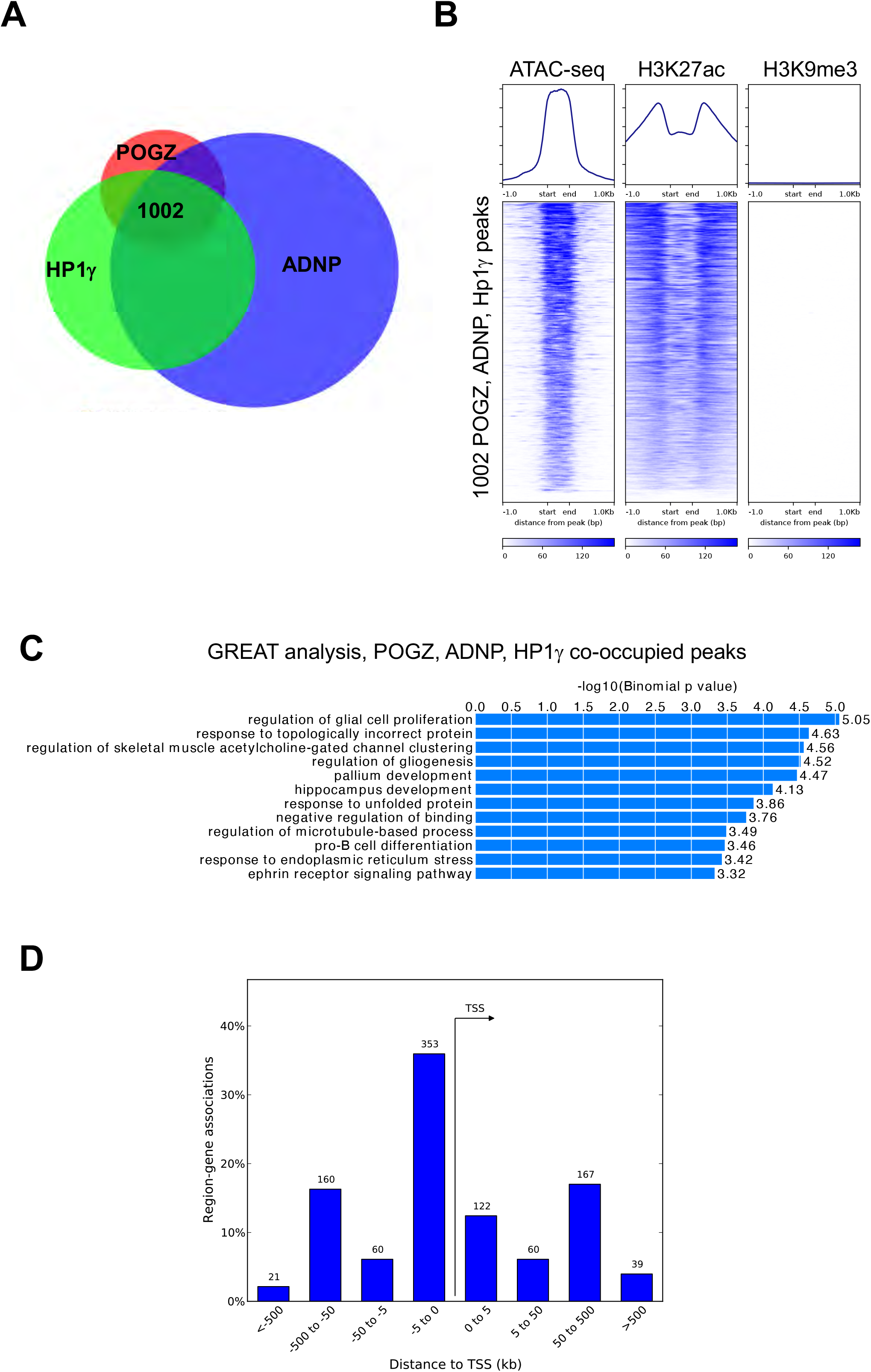
**S7A)** Venn Diagram of C&R peak overlaps from ADNP, HP1γ, and POGZ consensus peaks across replicates. S7B) Heat map of ATAC-seq, H3K27ac ChIP-seq, and H3K9me3 ChIP-seq signal across 1002 loci co-occupied by POGZ, ADNP, and HP1γ in E13.5 telencephalon. S7C) GO analysis of nearest genes to loci co-occupied by POGZ, ADNP, and HP1γ. S7D) POGZ, ADNP, and HP1γ co-occupied C&R peaks linked to nearest genes, binned by distance and orientation from gene TSS using GREAT.

**Figure S8.**
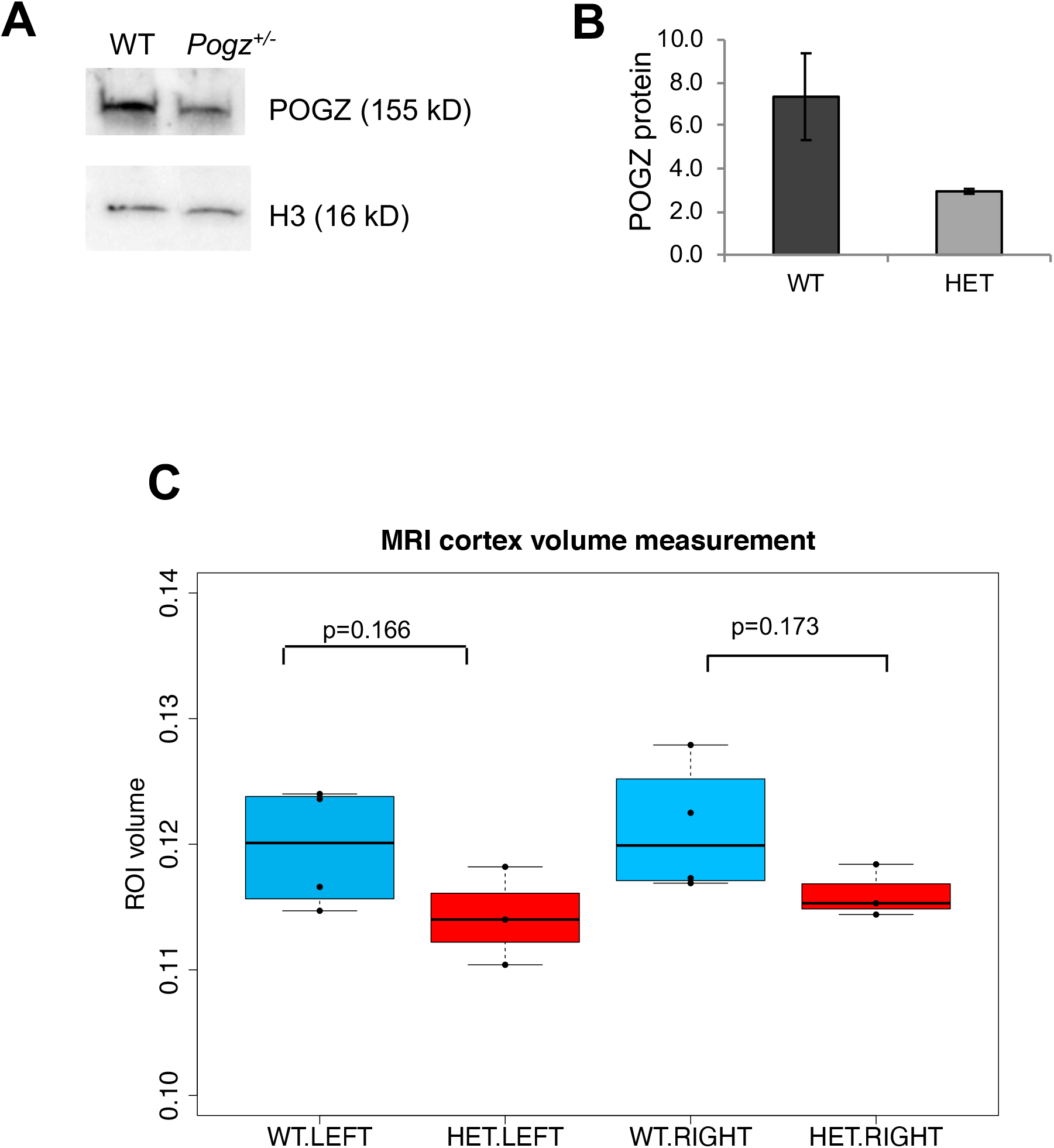
**S8A)** Western blot using anti-POGZ antibody in nuclear extracts from P2 cortex of *Pogz^+/−^* and wildtype. Histone H3 loading control. S8B) Quantification of anti-POGZ signal from multiple Western blots from P28 *Pogz^+/−^* and wildtype, normalized to loading control. S8C) MRI quantification of left and right hemisphere cortex volume from P28 *Pogz^+/−^* and wildtype mice.

## METHODS

### RESOURCE AVAILABILITY

Further information and request for resources should be directed to the lead contact, John Rubenstein.

#### Materials Availability

All unique reagents generated in this study are available from the lead contact with a completed Materials Transfer Agreement

#### Data Availability

RNA-seq, Histone ChIP-seq, ATAC-seq, and CUT&RUN data will be deposited to GEO.

### EXPERIMENTAL MODEL AND SUBJECT DETAILS

#### Animal Models

All procedures and animal care were approved and performed in accordance with National Institutes of Health and the University of California San Francisco Laboratory Animal Research Center (LARC) guidelines, UCSF IACUC approval number AN180174-02.

#### Developing Human Brain Samples

Developing human brain samples were obtained with patient consent in strict observance of the legal and institutional ethical regulations. Protocols were approved by the Human Gamete, Embryo, and Stem Cell Research Committee, and the Institutional Review Board at the University of California, San Francisco. Fresh fetal brain samples were obtained from elective terminations, with no karyotype abnormalities or genetic conditions reported, and transported in freshly made Cerebral Spinal Fluid on ice (CSF). Samples ranged from 17gw to 18gw in age. All dissections and experiments were performed within two hours of tissue acquisition. Dissections of the cortical sample acquired included the entire telencephalic wall, from the ventricular zone to the meninges.

#### CRISPR mouse generation

SgRNAs were designed using the guide design tool at crispr.mit.edu. The following sgRNAs in Table S6 were cloned into the px330 vector (Zhang lab) and sgRNAs were generated by in vitro transcription using the MEGAshortscript T7 transcription kit (Invitrogen). Both sgRNAs were injected into fertilized mouse oocytes together with Cas9 protein at the Gladstone Transgenic Core facility. Mice were screened for 10 kb deletions using PCR Primers (Table S6) upstream and downstream of the two sgRNA target sites.

### METHOD DETAILS

#### *In situ* hybridization

E13.5 whole head was dissected and postfixed overnight in 4% paraformaldehyde, and transferred to 30% sucrose overnight. 20 micron thick cryosections were obtained. In situ hybridization using antisense RNA probes in Table S6 was performed as described (Sandberg et al. 2018). Sections were developed at 37C and were imaged two days later.

#### Immunofluorescence

E13.5 whole head was dissected and postfixed overnight in 4% paraformaldehyde, and transferred to 30% sucrose overnight. 20 micron thick cryosections were obtained. Immunostaining was performed using antibodies in Table S6 in 1x PBS 0.1% Triton and 4% donkey serum overnight at 4C. IF images were obtained using a Zeiss Confocal Microscope at 20x or 63x magnification. Cortical cell counts and layer thickness were quantified using ImageJ.

#### Luciferase Assay

Primers in Table S6 were used to amplify predicted regulatory elements bound by POGZ from mouse genomic DNA, then cloned into the minimal promoter pGl4.23 luciferase vector (Promega) using SacI and XhoI restriction sites (underlined) in the vector’s multiple cloning site.

Neonatal cortical and basal ganglia tissues were dissected from CD1 mice in cold EBSS, followed by trypsin (Thermo Fisher Scientific 25200056) treatment for 15 minutes at 37°C. Trypsinization was inhibited using 10% FBS containing DMEM. Cells were washed once with DMEM, then resuspended in 10% FBS containing Neuralbasal-A medium (Thermo Fisher Scientific 12348017) with B27 (Thermo Fisher Scientific 17504044). Cell density was quantified using hemocytometer. Cells were plated in poly-D-lysine and laminin coated coverslips (Corning 08-774-385) preloaded in 96 well plates and cultured in 37°C incubator for 14 days. Serum free Neuralbasal-A medium with B27 and Glutamax (Thermo Fisher Scientific 35050061) was used to maintain the cell growth.

Confluent cells were transfected in three 96-well plates with luciferase vectors (predicted enhancer-pGL4.23 or empty vector pGL4.23) and pRL renilla vector. Two days later, cells were lysed and luciferase levels detected using the Promega dual reporter luciferase assay kit. Luciferase levels were normalized to Renilla and averaged across three replicate experiments.

#### Co-Immunoprecipitation

Microdissected cortices from wild type E13.5 mouse embryos or 18 gw human embryos were triturated using P1000 tip 10 times to generate single cell suspension. To isolate nuclei, cells were resuspended in 1mL hypotonic lysis buffer (10mM HEPES pH 7.4, 10mM NaCl, 1.5mM MgCl_2_, 0.5mM DTT, and 1x Roche complete protease inhibitor) per 1E7 cells, held on ice for 10 minutes, and dounced 10-15 times with tight clearance pestle. Nuclei were collected by centrifugation and resuspended in nuclear extraction buffer (20mM HEPES pH 7.4, 300mM NaCl, 25% glycerol, 0.1mM EGTA, 0.02% Igepal, 0.5mM DTT, 250U Benzonase, and 1x Roche complete protease inhibitor) and incubated at 37°C for 15 minutes to digest DNA and RNA. Nuclei were rotated for 30 minutes at 4°C and then centrifuged for 30 minutes at 14,000 x g at 4°C to pellet insoluble fraction. Nuclear extracts were diluted 1:1 with nuclear extraction buffer lacking NaCl for a final concentration of 150mM.

For immunoprecipitations, Dynabeads Protein A (Thermo Fisher, 10001D) were washed twice with IP buffer and added to 5-10μL of antibody suspended in 1% BSA in PBS for two hours at room temperature. Beads were washed twice more with IP buffer and added to nuclear extracts overnight at 4°C. After separation from unbound lysate, beads were washed 5 times with IP buffer and bound proteins were eluted by incubating in 100μL Laemmli Sample Buffer at 70°C for 10 minutes.

#### Western Blot

Nuclear extracts were prepared as described above for Co-Immunoprecipitation, from wildtype, heterozygote, and homozygote cortex. 1ug protein was loaded for each genotype in 2x Laemmli Sample Buffer, and POGZ was blotted using the recommended dilution of antibody (1:1000); anti Histone H3 antibody was included for loading control. Relative levels of POGZ protein were measured by pixel intensity analysis of scanned Western blot images using ImageJ.

#### MRI

4% PFA-fixed brains were washed twice in 20 mL PBS for a total of 24 hours and imaged in Fluorinert FC-40 (Sigma Aldrich) for null background signal. The imaging was done on a 600 MHz NMR spectrometer (Agilent Technologies Inc.) with imaging gradients and the following parameters: 3D gradient echo, TE/TR 15/75 ms, 8 averages, field of view (FOV) 12.8 mm isotropic, resolution of 50 μm × 50 μm × 100 μm, and a total scan time of 5.5 hours. The acquired images were converted on console to the DICOM format. Volumetric measurements were made using Horos, an open source image viewer.

#### RNA-seq

From each dissection of E13.5 cortex or basal ganglia, RNA was isolated using QIAGEN RNeasy mini columns. RNA was treated with Turbo DNAse and inactivated. RNA quality was assessed using Agilent Bioanalyzer RNA Nano kit, and samples with RNA Integrity Number values greater than 9.0 were used for subsequent profiling. RNA-seq libraries were generated using Nugen’s Ovation Mouse RNA-seq kit and amplified for 15 cycles. Libraries were quantified using Agilent Bioanalyzer High Sensitivity DNA kit, and sequenced on Hiseq 2500 using paired end sequencing.

#### ATAC-seq

From each dissection of E13.5 cortex or basal ganglia, intact nuclei were isolated by pipetting up and down twenty times in ice cold 0.5 mL Buffer 1 (300mM sucrose, 60mM KCl, 15mM NaCl, 15mM Tris-HCl, pH 7.5, 5mM MgCl2, 0.1mM EGTA, 1mM DTT, 1.1mM PMSF, Protease inhibitors), and then lysed on ice for 10 minutes after adding 0.5 mL Buffer 2 (300mM sucrose, 60mM KCl, 15mM NaCl, 15mM Tris-HCl, pH 7.5, 5mM MgCl2, 0.1mM EGTA, 0.1% NP-40, 1mM DTT, 1.1mM PMSF, Protease inhibitors). During these ten minutes, nuclei were counted using trypan blue and 50,000 nuclei were spun down at 7,000rpm for ten minutes at 4C. Nuclei were resuspended in 25uL Tagmentation buffer, 22.5 uL Nuclease Free H20, and 2.5 uL Tagmentation Enzyme from Nextera DNA Library Prep Kit, gently mixed, and placed in 37C water bath for thirty minutes. The tagmentation reaction was stopped by MinElute PCR purification and DNA was eluted in 10uL Nuclease Free water. ATAC-seq library generation was performed using Illumina barcode oligos as described (Buenrostro et al 2015), for 8-11 cycles PCR using NEBNext High Fidelity 2x PCR master mix. The number of cycles was empirically determined for each library by qPCR. Libraries were bioanalyzed using Agilent High Sensitivity DNA Kit, pooled together and sequenced on Hiseq 2500 using paired end sequencing.

#### ChIP-seq

From each dissection of E13.5 cortex, intact nuclei were isolated by pipetting up and down twenty times in ice cold 0.5 mL Buffer 1 (300mM sucrose, 60mM KCl, 15mM NaCl, 15mM Tris-HCl, pH 7.5, 5mM MgCl2, 0.1mM EGTA, 1mM DTT, 1.1mM PMSF, 50mM Sodium Butyrate, EDTA-free Protease inhibitors), and then lysed on ice for 10 minutes after adding 0.5 mL Buffer 2 (300mM sucrose, 60mM KCl, 15mM NaCl, 15mM Tris-HCl, pH 7.5, 5mM MgCl2, 0.1mM EGTA, 0.1% NP-40, 1mM DTT, 1.1mM PMSF, 50mM Sodium Butyrate, EDTA-free Protease inhibitors). During this ten minutes, nuclei were counted using trypan blue and 500,000 nuclei were spun down at 7,000rpm for ten minutes at 4C. Nuclei were resuspended in 0.250mL MNase buffer (320mM sucrose, 50mM Tris-HCl, pH 7.5, 4mM MgCl2, 1mM CaCl2, 1.1mM PMSF, 50mM Sodium Butyrate) and incubated in a 37C water bath with 2 microliters Micrococcal Nuclease enzyme (NEB) for eight minutes. Micrococcal Nuclease digestion was stopped by adding 10 microliters 0.5M EDTA, and chromatin was spun down for 10 minutes 10,000rpm 4C. Soluble fraction “S1” supernatant was saved at 4C overnight, and “S2” fraction was dialyzed overnight in 250uL dialysis buffer at 4C (1mm Tris-HCl pH 7.5, 0.2mM EDTA, 0.1mM PMSF, 50mM Sodium Butyrate, Protease Inhibitors). Next day S1 and S2 fractions were combined, 50 microliters were saved as input, and Chromatin immunoprecipitation was set up in ChIP buffer: 50mM Tris, pH 7.5, 10mM EDTA, 125 mM NaCl1, 0.1% Tween. 250m M Sodium Butyrate was supplemented for H3K27ac ChIPs. 1 microliter of antibody was added to 1mL chromatin in ChIP buffer and incubated overnight at 4C rotating. Protein A and Protein G beads (10 microliters for each ChIP) were blocked overnight in 700uL ChIP buffer, 20 uL yeast tRNA (20mg/mL), and 300uL BSA (10mg/mL). Beads were washed three times for five minutes on ice in Wash buffer 1 (50 mM Tris, pH 7.5, 10mM EDTA, 125mM NaCl, 0.1% Tween-20, with protease inhibitors and 5mM sodium butyrate) and three times in Wash buffer 2 (50 mM Tris, pH 7.5, 10mM EDTA, 175mM NaCl, 0.1% NP-40, with protease inhibitors and 5mM sodium butyrate), and ChIP DNA was eluted in elution buffer at 37C and purified by phenol chloroform extraction and ethanol precipitation. Sequencing libraries were made using Nugen Ovation Ultralow V2 kit and quantified by Agilent High Sensitivity DNA kit on the Agilent bioanalyzer.

#### CUT&RUN

From each dissection of E13.5 telencephalon, intact nuclei were isolated using Buffer 1 and Buffer 2 as for ChIP-seq and ATAC-seq above. From each dissection of human cortical tissue, intact nuclei were isolated by douncing in 1mL Buffer 1 with loose pestle and lysing in 1mL Buffer 2. After spinning down nuclei at 7,000rpm for ten minutes at 4C, we methodically followed the protocol for CUT &RUN (Skene, Henikoff, and Henikoff 2018), starting at step 6 “Resuspend in 1mL of wash buffer at RT by gentle pipetting”. Antibodies were used at the following dilutions: POGZ 1:500, ADNP 1:13, Hp1γ 1:500, IgG 1:1000. For the final DNA extraction step, we performed phenol-chloroform extraction and ethanol precipitation. We generated libraries using Nugen’s Ovation Ultralow V2 kit, and amplified libraries for 16 cycles, and bioanalyzed libraries using the Agilent High Sensitivity DNA kit and sequenced libraries on the Hiseq 2500.

#### ATAC-seq analysis

We mapped groomed fastq files to the mm9 genome using Bowtie2 default mode (Langmead and Salzberg 2012). Samtools merge (v1.10) was used to merge experimental replicates together before peak calling (Li et al. 2009). For peak calling, we used MACS2 (v2.1.1) *callpeak* (Y. Zhang et al. 2008). We disabled model-based peak calling and local significance testing. We used a fixed fragment extension length of 200bps, and we used a q-value cut-off of 0.05. We used the peak calling results to run MACS2 differential peak calling (bdgdiff) with a min peak length of 150bps. Relative ATAC-seq signal in wildtype versus *Pogz^−/−^* was generated using DeepTools bamCompare tool, dividing the genome into bins of 50 kb.

#### ChIP-seq analysis

We mapped groomed fastq files to the mm9 genome using Bowtie2 default mode (Langmead and Salzberg 2012). We used samtools merge (v1.10)(Li et al. 2009) to merge experimental replicates together before peak calling. For peak calling, we used MACS2 (v2.1.1) *callpeak* (Y. Zhang et al. 2008). We disabled model-based peak calling and local significance testing. We used a fixed fragment extension length of 200bps, and we used a q-value cut-off of 0.01. We used the peak calling results to run MACS2 differential peak calling (bdgdiff) with a min peak length of 150bps.

#### Motif analysis

We used Homer to call motifs on our peak sets, using a size window of 200 (Heinz et al. 2010). We used the findMotifsGenome.pl and annotatePeaks.pl tools to acquire sets of enriched motifs for all our experiments and annotations for our peaks. We used MEME-Chip to call *de novo* motifs in our peak sets, using default settings (Bailey et al. 2015). We subsequently used FIMO with default settings to scan POGZ C&R peaks for individual instances of *de novo* motifs (Bailey et al. 2015). From FIMO output we determined if the instance of the motif was degenerate or matched the consensus motif exactly (non-degenerate), and we generated heatmaps of ATAC-seq on those peak sets using DeepTools computeMatrix and plotHeatmap tools (Ramírez et al. 2016).

#### CUT&RUN analysis

We mapped groomed fastq files to the mm9 genome using Bowtie2 default mode (Langmead and Salzberg 2012). Samtools merge (v1.10) was used to merge experimental replicates together before peak calling (Li et al. 2009). For peak calling, we used MACS2 (v2.1.1) *callpeak* (Y. Zhang et al. 2008). We disabled model-based peak calling and local significance testing. We used broad peak settings and a q-value cut-off of 0.05. Bedtools (Quinlan and Hall 2010) was used to determine consensus peaks across replicates and antibodies. Heatmaps of CUT&RUN signal for various antibodies over POGZ C&R peaks were generated using DeepTools computeMatrix and plotHeatmap tools (Ramírez et al. 2016). To plot CUT&RUN signal at consensus POGZ/ADNP/HP1γ peaks proximal to differentially expressed genes, we used Bedtools to intersect consensus peak bed files with bed files of differentially expressed genes (qvalue<0.01) from cortex *Pogz^−/−^* (Table S2) extended by 100kB upstream and downstream of gene locations. We then used DeepTools computeMatrix and plotHeatmap tools (Ramírez et al. 2016) to plot the distribution of CUT&RUN signal for POGZ, ADNP, and HP1γ at those peaks.

#### RNA-seq analysis

RNA-seq reads were mapped to mm9 using HISAT2 version 2.0.5 (Kim et al. 2019), and reads were counted on mouse genes using htseq version 0.6.1p1 (Anders, Pyl, and Huber 2014). Differentially expressed transcripts were determined from count tables using DESeq2 version 1.14.1, and genes with qvalue<0.05 were included in plots and tables of differentially expressed genes. For RNA-seq analysis of transposable element expression, we mapped reads to the mm9 genome using RNA STAR (Dobin et al. 2013) and counted reads on repeatmasked sequences of the mm9 genome using featureCounts (Liao, Smyth, and Shi 2014). Differentially expressed transcripts were determined from count tables using DESeq2, and genes with qvalue<0.05 were included in MA plots and tables.

#### GO Analysis

GO analysis for genomic loci (CUT&RUN, differential ATAC-seq, H3K27ac ChIP-seq peaks) was performed using GREAT version 4.0.4 (McLean et al. 2010) and the single nearest gene setting for associating genomic regions with genes. GO analysis for differentially expressed genes (RNA-seq) was performed using DAVID Bioinformatics Resources 6.8 (Huang, Sherman, and Lempicki 2009).

#### Transposable Element Analysis

Performed separately for human (hg19) and mouse (mm9) genomes, each genome was divided into non-overlapping 100bp windows (excluding ENCODE blacklisted regions) and intersected with repeat coordinates from Tandem Repeats Finder

(obtained from the UCSC Genome Browser) as well as POGZ peaks (CUT&RUN). This was performed for each repeat family using separate euchromatic and heterochromatic POGZ peaks. The odds ratio and p-value was computed from the contingency table of genomic bins overlapping a particular repeat family versus those overlapping a particular set of CUT&RUN peaks. P-values were adjusted for multiple testing (Benjamini-Hochberg).

#### Gene set enrichment analysis

We tested if various gene sets (ASD risk genes, Developmental Delay risk genes, CHD8 targets, Fragile-X targets, lung expressed genes, liver expressed genes, olfactory receptor genes, and random subsets of 512 genes expressed with mean rpkm>3 in whole brain from Brainspan 16gw-19gw samples) were enriched near two different sets of C&R peaks (POGZ and H4K20me3). Enrichment was tested over the space of all protein coding genes, with one vector representing if a gene was in the gene set, and another vector representing if a gene was within 50 kilobases of a statistically significant C&R peak. A 2×2 contingency table was computed from these vectors, and Fisher’s exact test used to compute the odds ratio and p-value. The resulting p-values from all gene set and C&R target combinations were corrected for multiple testing (Benjamini-Hochberg).

We also tested if there was a statistically significant difference in the proportion of CHD8 target genes proximal to POGZ peaks versus the proportion of whole brain expressed genes proximal to POGZ peaks. The latter proportion was computed using 1000 different random samples of expressed whole brain genes, matched in size to the CHD8 target gene set, and the empirical p-value of the proportion of POGZ-proximal CHD8 target genes was computed using the resampled distribution of POGZ-proximal expressed whole brain gene proportions. This resulted in an empirical p-value of 0, as the maximum proportion of POGZ-proximal random whole brain genes was 56.9 (mean 53.4), while the proportion of POGZ-proximal CHD8 gene targets was 68.2.

All genes names were converted to their current HUGO names for compatibility. Whole brain genes with mean RPKM > 3 for 16-19 gestational week samples in RNA-seq data from Brainspan were defined as expressed.

## Acknowledgements

This work was supported by the research grants to JLR from: Nina Ireland, The Simons Foundation, NINDS R01 Ns099099. Duan Xu acknowledge grant R35NS097299.

J.L.R. is cofounder, stockholder, and currently on the scientific board of *Neurona*, a company studying the potential therapeutic use of interneuron transplantation.

## Notes

### Competing Interest Statement

The authors have declared no competing interest.

